# Improved methodology for protein NMR structure calculation using hydrogen bond restraints and ANSURR validation: the SH2 domain of SH2B1

**DOI:** 10.1101/2022.08.29.505644

**Authors:** Nicholas J. Fowler, Marym F. Albalwi, Subin Lee, Andrea M. Hounslow, Mike P. Williamson

## Abstract

Protein structures calculated using NMR data are less accurate and less well defined than they could be. Here we use the program ANSURR to show that this deficiency is at least in part due to a lack of hydrogen bond restraints. We then describe a protocol to introduce hydrogen bond restraints into the structure calculation of the SH2 domain from SH2B1 in a systematic and transparent way, and show that the structures generated are more accurate and better defined as a result. We also show that ANSURR can be used as a guide to know when the structure calculation is good enough to stop.

## Introduction

We recently introduced a new method, ANSURR (Accuracy of NMR structures Using RCI and Rigidity), which permits a simple and robust calculation of the accuracy of NMR structures.^1-3^ We noted^2^ that when estimated in this way the accuracy of NMR protein structures improved until about 2005, but has not changed much since that point; and we suggested that one reason for the lack of improvement since then is that there has been until now no reliable way to calculate accuracy, and therefore no way to measure the success of technical improvements. With the introduction of ANSURR, we are now able to measure whether new methods are effective. In this paper, we address one experimental restraint that has over the years proved difficult to implement, namely hydrogen bond restraints.

The most widely used structural restraints for NMR structure calculation are nuclear Overhauser effects (NOEs). These are easily obtained from NOESY spectra, and have (in principle) a simple interpretation, in that under appropriate conditions, an NOE peak denotes the close distance between two protons, with an intensity proportional to *r*^-6^, where *r* is the distance between the two nuclei.^4^ Another widely used restraint is to use backbone chemical shifts to produce restraints on backbone dihedral angles, using a database approach, for example using the program TALOS-N.^5^

Hydrogen bonds are one of the few other common restraints, and are very strong restraints. They are typically implemented as a pair of distance restraints: one from the hydrogen donor to its acceptor (a carbonyl oxygen in the case of a backbone-backbone hydrogen bond), and one between their attached heavy atoms (amide nitrogen to carbonyl carbon). These two distances can be used in combination to help to restrict hydrogen bond geometries appropriately. It is a strong restraint because the distances are short and have a small range between upper and lower bounds. Violations of the restraint therefore typically become energetically unfavourable with relatively small changes in geometry. For this reason, a hydrogen bond restraint (HBR) should be applied with caution, because it effectively forces the relevant groups to form a hydrogen bond.

Hydrogen bonds are more difficult restraints to apply than NOEs because they cannot usually be obtained directly from the experimental data. NOE restraints come directly from NOESY spectra, and dihedral restraints come more or less directly from backbone chemical shifts. It is possible to observe hydrogen bonds directly from observation of three-bond scalar couplings across the hydrogen bond.^6^ However, the coupling constant is very small and for most proteins the resultant signal is too weak to be detected. Therefore the presence of a hydrogen bond has to be inferred. Hydrogen bonds in regular secondary structure can be predicted based on NOE patterns in the vicinity,^7^ but a more secure identification of hydrogen bonds relies on secondary factors such as slow amide exchange of the hydrogen bonded amide proton^8-9^ or on the magnitude of the amide proton temperature coefficient.^10^ Neither of these is a completely reliable guide, meaning that any inference of a hydrogen bond is potentially wrong. Significantly, amide exchange and temperature coefficients identify the amide group but do not identify the carbonyl donor. Therefore for most hydrogen bonds, identification of the donor is based on inferences rather than on direct observation.

For these reasons, NMR spectroscopists have generally been careful in how they apply HBRs in protein structure calculation. There is no “standard protocol” but typically HBRs are only applied towards the end of the structure calculation, as a way of tidying up the structure, the suspicion being that it may not be scientifically justified to use them as part of the standard structure calculation. There has thus emerged a sense that HBRs are not as “respectable” as NOEs or dihedral restraints, as a result of which, the details of hydrogen bond restraints are often deeply buried in papers and it is often very hard to work out from the literature exactly how HBRs were generated and used. This is a pity, because HBRs are such powerful restraints, and therefore very useful if used correctly. The main aim of this work is to propose a scientifically justifiable and transparent protocol. We describe a robust iterative approach to the addition of HBRs, monitored using ANSURR, and demonstrate the effect of HBRs on the structure calculation of the SH2 domain from the scaffold protein SH2B1.

SH2B1 is a signalling scaffold protein that acts downstream of the insulin and leptin receptors^11^ and regulates the activity of the associated JAK2 kinase. Mutations to the SH2 domain of SH2B1 have been shown to disrupt the leptin signalling pathway;^12^ deletion leads to an obese phenotype and severely disrupts signalling downstream of the insulin receptor.^13^ SH2B1 consists of a dimerization domain, a PH domain and an SH2 domain, which together form less than half of the protein, the rest being intrinsically disordered.^11^ Part of the function of SH2 is to bind to JAK2 kinase via its phosphorylated pTyr^813^ and modulate its kinase activity.^14^ It is thus a protein with important biological effects. It is also an example of a well-studied domain with a crystal structure. For the purposes of this study this is useful because the crystal structure provides us with a useful guide to the accuracy of our calculations.

## Experimental Section

### A set of comparable NMR and X-ray structures

The Structure Integration with Function, Taxonomy and Sequence (SIFTS) resource^15^ was used to match NMR structures to X-ray structures. NMR structures were required to have at least 80% backbone chemical shift completeness. X-ray structures were selected based on those which gave most coverage of the NMR structure, then which of those had the best resolution. Only pairs where the X-ray structure covered at least 80% of the NMR structure were used. Redundancy was reduced by randomly selecting a single structure pair for each UniProt accession number (resulting in 222 pairs) before applying a 50% sequence identity cut-off, leaving 215 structure pairs for analysis. A list of paired NMR and X-ray structures is provided in Table S1.

### Protein expression and purification

The gene for SH2 was made synthetically (Integrated DNA Technologies). The gene sequence is that of mouse SH2 (Uniprot Q91ZM2), codon optimised for expression in *E. coli* (Integrated DNA Technologies). The synthetic gene was 461 bp long and consisted of N-terminal homology region, ribosome binding site, start codon, His tag, protein sequence (118 amino acids), stop codon and C-terminal homology region. It was inserted into the pET-2 plasmid, which had been linearised by cutting with BamH1 and Xba1, following standard Gibson protocol.^16^ The plasmid was transformed into BL21 (DE3) *E. coli*, and grown in labelled M9 medium supplemented with 100 μg/ml ampicillin. It was incubated, induced with 0.5 mM IPTG, and grown overnight at 25°C. Cells were lysed by sonication and protein was purified using a Ni-NTA column, eluting with 300 mM imidazole, followed by polishing using a Superdex 200 column. For NMR, buffer was exchanged to 50 mM potassium phosphate pH 6, containing 1 mM trimethylsilyl propionate (TSP) as an internal standard, in 10% D_2_O. The protein concentration was approximately 1 mM in a Shigemi tube.

### NMR assignment

NMR data were collected on a Bruker DRX-600 equipped with a cryoprobe. Manual assignment was done using the program Asstools,^17^ which is based on a Monte-Carlo simulated annealing method. Three backbone amides in the HSQC spectrum have low intensity (Trp17, Gln61 and His81). The manual backbone assignment was complete except for Gly20, which appeared to be absent, and the two N-terminal residues. The final sequential assignment of SH2 backbone resonances showed that in total 95% of backbone resonances for non-proline residues 9 to 118 were assigned (96% of ^1^H, 96% of ^15^N, 96% of Cα, 94% of Cβ, 92 % of Cʹ), the missing signals being due to Trp17, Gly20, Gln61, and His81. Most sidechains were assigned, with the exceptions of some ends of sidechains, Phe42, Ile99, and Val115.

The automated chemical shift assignment was performed in two steps, following the recommended procedure:^18^ an ensemble of chemical shift assignments was computed followed by combining the resultant raw chemical shift assignments into a single consensus resonance list. The ensemble of chemical shift assignments was obtained from 20 runs of the GARANT algorithm, in which the iteration size for one generation was 100. Each independent run started from the same experimental peak lists but using a different random seed value, and optimised the match between observed peaks and expected peaks based on the knowledge of the amino acid sequence and the magnetization transfer pathways in the spectra used.^19^ The matching was done with the recommended tolerance values of 0.03 and 0.4 ppm for the ^1^H dimension and for the ^13^C and ^15^N dimensions, respectively.

### Structure restraints and calculations

Calculations were carried out using standard procedures in CYANA 3.98.5. Tolerance values for the chemical shift matching were 0.04 ppm for ^1^H dimension, 0.03 ppm for ^15^N or ^13^C bound ^1^H dimension, and 0.45 ppm for ^15^N and ^13^C dimensions. ^15^N NOESY, ^13^C NOESY for aliphatic atoms, and ^13^C NOESY for aromatic atoms (mixing times 100 ms) were sorted in XEASY format which contains peak positions and volumes. The input to CYANA was a list of manually checked assignments plus NOE pick lists, dihedral restraints and HBRs. Dihedral angle restraints (φ and ψ) were generated according to the backbone chemical shifts in the SH2 protein using the TALOS-N program.^5^ 93 pairs of TALOS-N angles that were predicted as strong or generous were converted into torsion angle restraints using a CYANA macro. No χ_1_ restraints were used. Error values were given a default value of twice the standard deviation listed by TALOS-N. The predicted dihedral angles of proline residues were excluded from the CYANA torsion angles list, even for those classified as strong, namely 98, 100 and 116. The calculation started with 100 conformers generated from random torsion angle values. Each conformer is generated after 10000 torsion angle dynamics steps. For each HBR iteration, CYANA was run with different random seeds, selecting the 20 structures with the lowest target function out of a set of 100. This was duplicated a total of 60 times at each iteration starting from the same NOE peak lists, implying that the NOE restraints assigned will be different for each of these duplicate calculations. At each HBR iteration, the HBRs generated from the previous iteration were applied as starting restraints. For the final iteration of CYANA calculations, the 60 different sets of NOE restraints from the previous iteration were analysed, and restraints were adopted as consensus restraints if they were identified in 36 out of the 60 sets, using the largest upper bound distance from the set. This resulted in a reduction in the total number of NOE restraints used, from roughly 1800 to 1344.

Subsequent refinements in explicit solvent were carried out in CNS^20^ using the protein-allhdg parameters. CYANA distance restraints were converted into upper distance restraints in CNS format, and TALOS-N dihedral restraints were regenerated in CNS format.

Additional structure restraints were obtained from temperature coefficients and from amide exchange. For temperature coefficients, HSQC spectra were obtained at 288, 293, 298, and 303 K. All chemical shifts were referenced to TSP at 0 ppm. Coefficients were obtained by least squares fitting of a straight line. Amides defined as potential hydrogen bond donors are those with fitted temperature coefficients of -4.5 ppb/K or greater.^10, 21^ Amide exchange rates were measured by lyophilising the protein from a solution at pH 5.5, redissolving in D_2_O, and running a series of HSQC spectra at 288 K. Slowly exchanging amides are those visible in HSQC spectra started within 9 minutes of dissolution. A full list of amides used can be found in Table S3.

Residual dipolar coupling restraints (RDCs) were obtained using compressed polyacrylamide gels. Gels were poured and set in open-ended 5 mm NMR tubes with an inner diameter of approximately 4.2 mm, using 7% acrylamide (an 18:1 w/w mixture of acrylamide to N,N’-methylenebisacrylamide) and polymerised using 0.1% w/v ammonium persulfate and 0.1% v/v N,N,N’,N’-tetramethyethylenediamine (TEMED).^22^ Gels were left overnight to set, pushed out of the tube, washed extensively in distilled water, and dried in an oven at 37 °C under a glass cover until they had shrunk to roughly one half of their original dimensions. They were then inserted into a Shigemi tube in the presence of 380 μM protein in 50 mM phosphate, pH 6, 10% D_2_O, and the Shigemi plunger was inserted so that the swelled gel would be 45% compressed relative to its initial height. RDCs were measured using the Bruker IPAP pulse program hsqcf3gpiaphwg.2,^23^ and processed to separate the two peaks into two different spectra, using zero filling in the ^15^N dimension to achieve 0.5 Hz/pt digital resolution. The estimated accuracy of measurement of RDCs is ± 1 Hz, and RDCs were in the range of -22 to +27 Hz. RDC *Q* factors were calculated using PALES.^24^

### Structure predictions using AlphaFold and RosettaFold

AlphaFold predictions were obtained using ColabFold v1.2^25^ (https://colab.research.google.com/github/sokrypton/ColabFold/blob/v1.2.0/AlphaFold2.ipynb) with the following options: MSA mode: MMseqs2 (UniRef+Environmental); use templates: yes; number of recycles: 3; refine with amber: yes. RosettaFold predictions were obtained using the Robetta web server^26^ (https://robetta.bakerlab.org/) with the default options.

### Other calculations

All ANSURR calculations were performed with ANSURR v1.2.1 (DOI 10.5281/zenodo.5655244) with the option to re-reference chemical shifts using PANAV. Ramachandran distributions were calculated using ramalyze, part of the Molprobity suite.^27^ Amide proton chemical shift calculations were made using SHIFTX2 1.10A ^28^. Hydrogen bonds in crystal and NMR structures (Fig. 1) were identified using FLEXOME [https://doi.org/10.15125/BATH-00940]^29^ using a cutoff of 0 kcal/mol for hydrogen bond energy.

**Figure 1.**
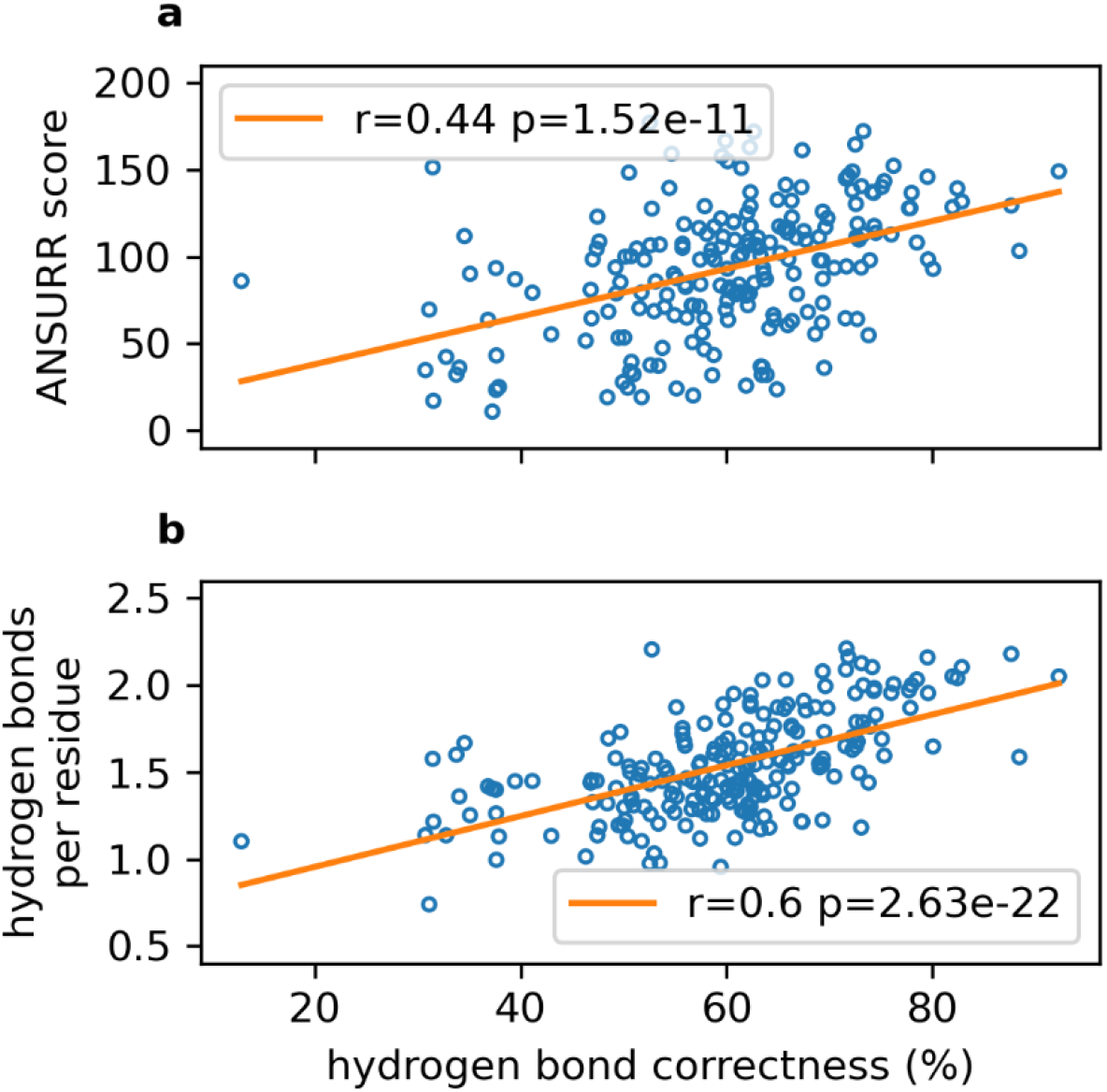
Correlations of structural parameters with hydrogen bond correctness, calculated as the average percentage of hydrogen bonds in an X-ray structure that appear in the corresponding NMR ensemble. (a) ANSURR score (b) number of hydrogen bonds per residue. Values were calculated for each structure in the ensemble and then averaged to give the value for the ensemble. The line of best fit is shown in orange, together with the Pearson correlation coefficient and two-tailed *p*-value.

## Results

### ANSURR suggests NMR structures are lacking hydrogen bonds

The aim of this study is to present an iterative approach to introducing hydrogen bond restraints (HBRs) into an NMR structure calculation, and to investigate whether this improves the accuracy of the structure. Our primary measure of accuracy is ANSURR. ANSURR calculates the rigidity of a protein structure using the algorithm FIRST^30^ and compares it to the rigidity derived from experimental backbone chemical shifts via the Random Coil Index (RCI).^31^ The similarity obtained by comparing these two measures provides the accuracy. The comparison uses two measures: *correlation*, which calculates the extent to which peaks and troughs in rigidity along the sequence match, and is primarily a measure of the correctness of local secondary structure; and *RMSD*, which compares rigidity along the sequence. Our previous studies^1-2^ showed that although the correlation statistic for NMR structures is often good, the RMSD is often poor, in almost all cases because NMR structures are too floppy in comparison to the rigidity indicated by backbone chemical shifts. We suggested that a major cause of this floppiness is that NMR structures have too few hydrogen bonds, in comparison to either crystal structures^2^ or AlphaFold structures^3^ of essentially identical proteins. This was the rationale behind the present study, which looks at the effect on accuracy of adding more HBRs. Before starting these structure calculations, we therefore undertook a more thorough study of the relationship between hydrogen bond content and accuracy.

A set of 215 pairs of published NMR and X-ray structures was assembled (Table S1). This set was used to compute the average hydrogen bond correctness of each NMR ensemble, defined as the average percentage of hydrogen bonds in an X-ray structure that appear in an NMR ensemble. This calculation relies on the assumption that the hydrogen bonds in a crystal structure are also present in solution, which is clearly not completely true, but is a good approximation.^1^ We computed the average ANSURR score (the sum of correlation and RMSD scores, and thus a ranked score out of 200) for each ensemble and compared this to the average hydrogen bond correctness.

Figure 1a shows that NMR structures with higher hydrogen bond correctness tend to be more accurate as measured by ANSURR, and also indicates that the hydrogen bonds in X-ray structures generally represent those that appear in solution. Plotting the number of hydrogen bonds per residue in each ensemble against hydrogen bond correctness (Figure 1b) demonstrates that poor scoring NMR ensembles are primarily inaccurate because they lack hydrogen bonds, rather than because they contain incorrect hydrogen bonds. We conclude that the accuracy of NMR structures will be improved by incorporation of HBRs, but only if they are correct. In the subsequent text, we explore how this can be achieved, monitoring the accuracy of the resultant structures using ANSURR and comparing to other available measures.

### NMR assignments for the SH2 domain and preliminary structure calculation

A plasmid for expression of mouse SH2 was constructed and inserted into *Escherichia coli*. The expressed protein is 118 residues long, with an 8-residue His-tag at the N-terminus, followed by the sequence D^9^QPLS…, with D^9^ corresponding to D^519^ of mature mouse SH2B, and finishes with the sequence YVPSQ. The protein was expressed and labelled uniformly with ^15^N and ^13^C. One aim of this project was to test out the automated assignment/structure calculation contained in FLYA/CYANA.^32^ The NMR spectrum was therefore assigned automatically using CYANA, as well as manually. The NMR spectra used are listed in Table S2, and the final manually assigned HSQC is shown in Figure S1. A comparison of the two assignments showed that 99% of the automated backbone chemical shift assignments (^1^H_N_, ^15^N, ^1^H_α_, ^13^C_ο_, ^13^C_α_ and ^13^C_β_) of SH2 (9-118) agreed with the manual assignment. The only resonances that were not assigned correctly by the automated calculation are the amide chemical shifts of Trp17 and Gln61, which are two of the four resonances that had missing signals (but assigned H and N) in the manual assignment. For aliphatic methyl and methylene sidechain groups, the agreement was 97%. On the other hand, many of the NOE-based chemical shift assignments did not match with the manual assignment. This includes the amide sidechains of Arg, Asn and Gln, and many aromatics: only 53% of these were assigned correctly. A graphic overview of the two assignments is shown in Figure S2. CYANA continues to check and refine its assignments during the course of structure calculations, and it is possible that some of these incorrect assignments would have been corrected. On the other hand, the accuracy of the automated structure calculation relies heavily on the accuracy of the first iteration of the structure calculation, which in turn depends on the accuracy of the initial set of assignments.^33^ The clear implication is that assignments should be checked manually when doing automated structure calculations.^34^ A manually checked set of assignments have been deposited in BioMagResBank^35^ under code 51342.

Structure calculation was carried out using the standard automated routines within CYANA. We used the default CYANA calculation, of 20 structures. We then repeated this calculation a total of 60 times changing the random seed each time, in order to obtain a better ‘consensus bundle’ of structures,^36^ giving a total of 1200 calculated structures per iteration. The reason for carrying out such a large number of calculations was to generate a reliable set of HBRs, with good discrimination between correct and incorrect HBRs. The figure of 60 was determined by calculating the ratio of true positive HBRs (that is, HBRs identified that are also present in the crystal structure) to false positive HBRs (HBRs identified that are not in the crystal structure). This ratio improved as the number of calculations was increased, reaching a plateau at 60 (Figure S3). HBRs were identified from these structures and used as restraints for the next iteration, as described below.

### Refining hydrogen bond networks using CYANA and sparse NMR data

The initial round of CYANA calculations produced an ensemble that clearly had the correct fold but was of low accuracy (as measured by ANSURR), and had poor convergence, a large RMSD between structures, and poor energies, all symptoms of a weak restraint set. We therefore improved the structures by iteratively and systematically including backbone hydrogen bond restraints, in such a way as to avoid incorrectly biasing the structure calculation (Figure 2). The protocol is structured to generate HBRs that are present in a significant fraction of ensemble structures, that are supported by experimental temperature coefficients and/or amide exchange rates, and that are consistent with other restraints. Iteration 6 was introduced as a way of removing any HBRs that were incompatible with the NOE and TALOS restraints.

**Figure 2.**
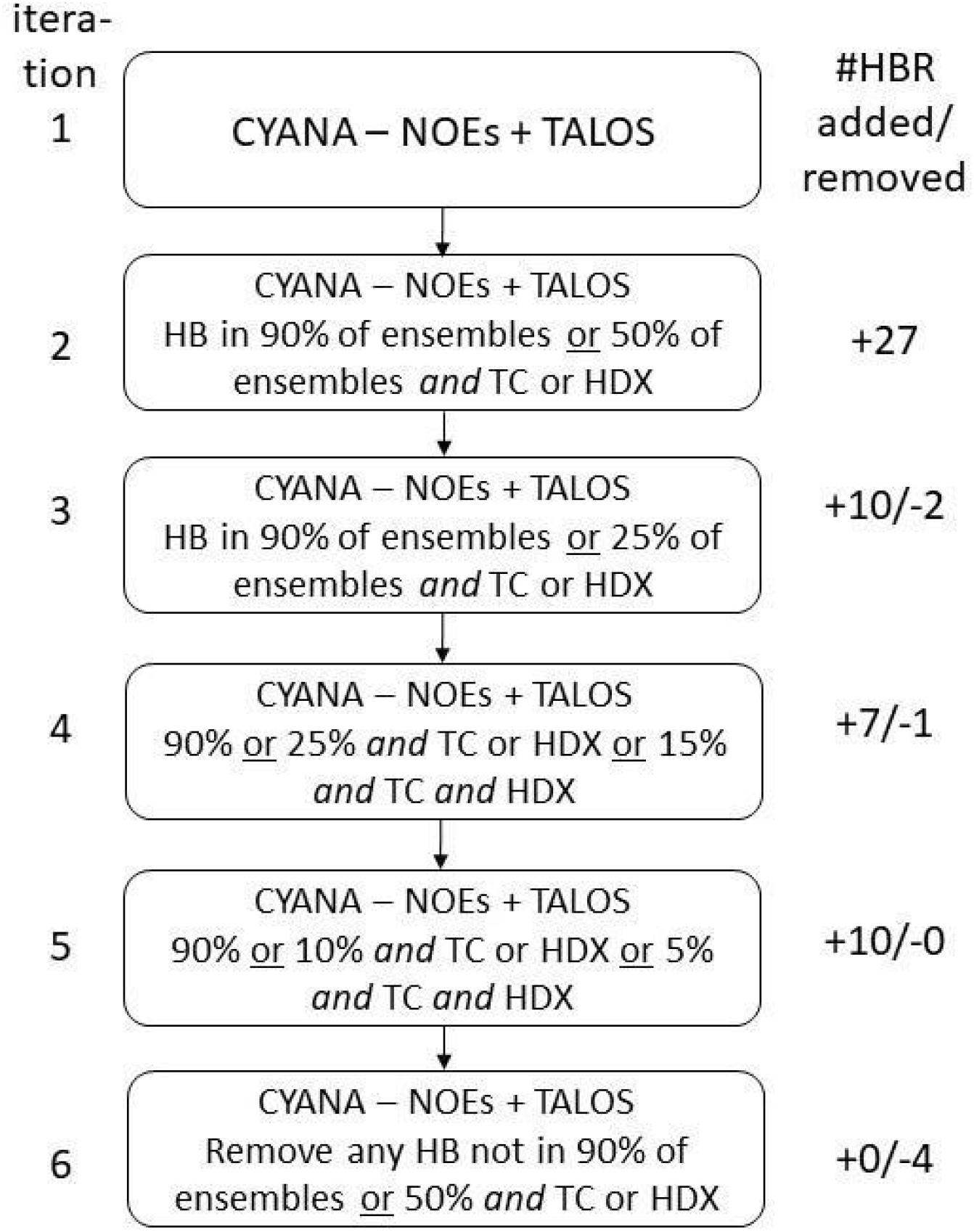
Protocol for addition of hydrogen bond restraints. 1200 structures (60 sets of 20) were calculated at each iteration. TC: temperature coefficient greater than -4.5 ppb/K. HDX: slow amide exchange. At each iteration, HBRs were added using the criteria listed, and calculations were repeated until there were no further changes in HBRs. Figures on the right are the number of hydrogen bonds added and removed at each iteration. In iteration 2, hydrogen bonds were used as restraints if either they appeared (ie, observed in at least 6 models within the ensemble of 20) in at least 90% of the ensembles generated in iteration 1, or appeared in at least 50% of the ensembles and were supported by either temperature coefficient or slow amide exchange data. In iteration 3 we adjusted our criteria so that we could accept less frequently sampled hydrogen bonds (25% of ensembles) so long as they were supported by temperature coefficient or slow amide exchange data. In iteration 4, we further lowered our acceptance criteria to explore rarely sampled hydrogen bonds that were supported by both amide exchange and temperature coefficients, with the view that the set of restraints already applied in more restrictive previous iterations should prevent the adoption of grossly incorrect hydrogen bond restraints at this stage. The criteria were relaxed further in iteration 5. Finally, for iteration 6 we restored our initial acceptance criteria to remove any hydrogen bond restraints that failed to converge, and were therefore potentially in conflict with NOE or dihedral restraints, or with accessible protein geometry.

The complete process identified 47 hydrogen bond restraints. Most of these occur in regular secondary structure. The final set of HBRs is illustrated in Figure 3. Not surprisingly, the first HBRs to be identified came from the helices and from the “core” β-strands (residues 41-69). The shorter β-sheet HBRs tended to be added later. Many of the HBRs that do not form part of regular secondary structures (for example, HBRs from the amide protons of G14, Y15, G104 and V109) were identified later. More details are available in Table S3.

**Figure 3.**
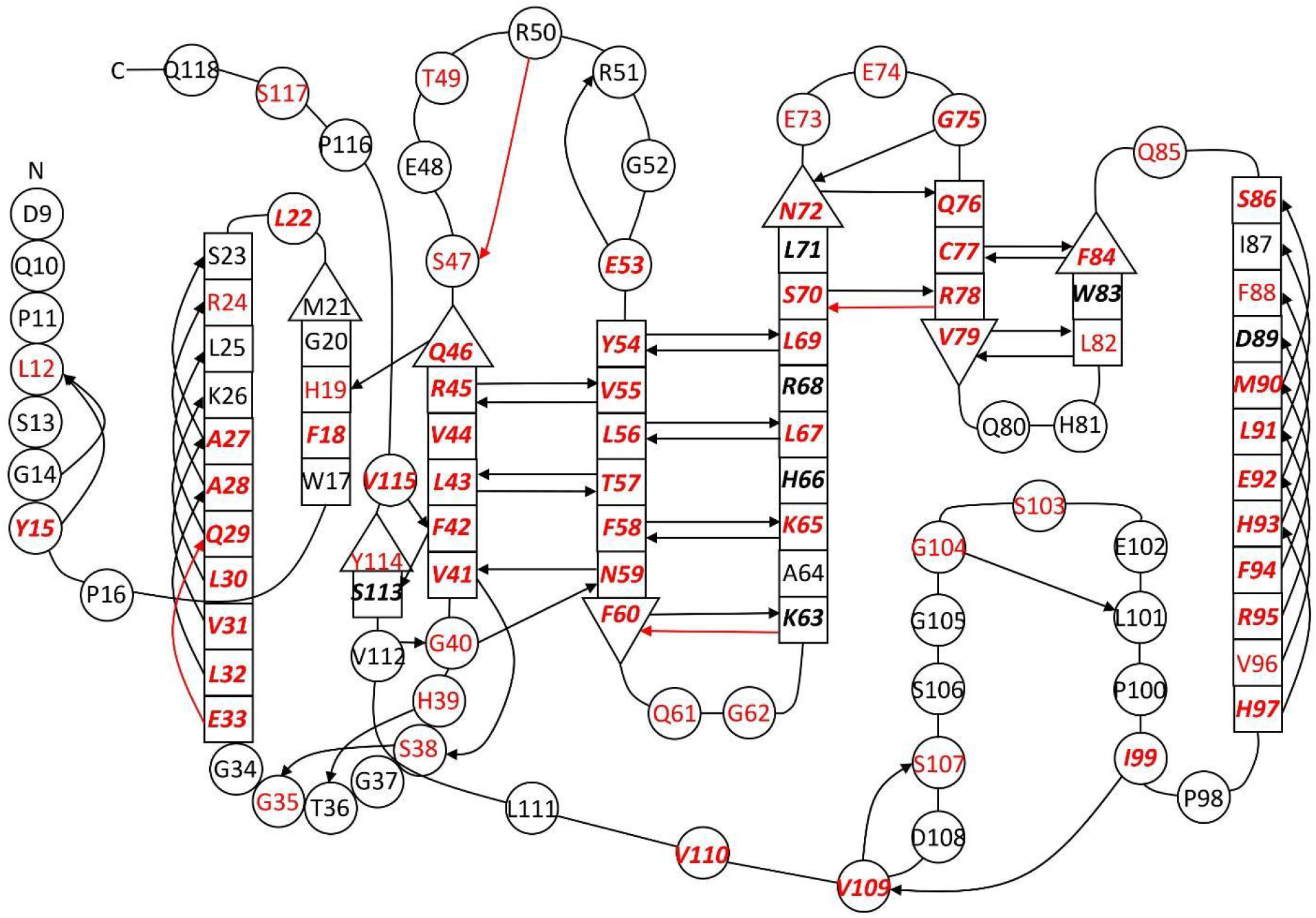
Secondary structure diagram for SH2. Residues in regular secondary structure are in square boxes, with an arrow at the C-terminal end of β-strands. Arrows indicate the hydrogen bond restraints used in the final iteration, starting at the NH and pointing towards the CO. Four restraints in red were used in iteration 5 and removed in the final iteration (Table S3). Residues are labelled in red if they have a temperature coefficient greater than -4.5 ppb/K, and in italic bold if they are slowly exchanging. We note that most residues with both slow exchange and large temperature coefficient have identified hydrogen bonding partners in our final iteration (42 out of 47).

At each stage in the calculation, the structures were analysed to measure whether the additional HBRs were improving the quality of the calculated structure ensemble; note that the structures shown here come directly from the CYANA calculation and have not yet been refined in explicit solvent. Our primary analytical tool was ANSURR. Figure 4a shows how ANSURR scores steadily increase as the number of HBRs is progressively increased, suggesting that inclusion of hydrogen bond restraints does improve structure accuracy. We note that the final iteration was designed to remove HBRs that do not have good supporting evidence. This iteration removed 4 out of 51 HBRs, leading to a drop in ANSURR score as well as some of the other measures of accuracy discussed below. The implication is that we probably removed correct HBRs, so this final step may have been over-cautious. The best scoring models in the final iteration 6 approach the accuracy of AlphaFold2^37^ and RosettaFold^26^ (Figure 4a).

**Figure 4.**
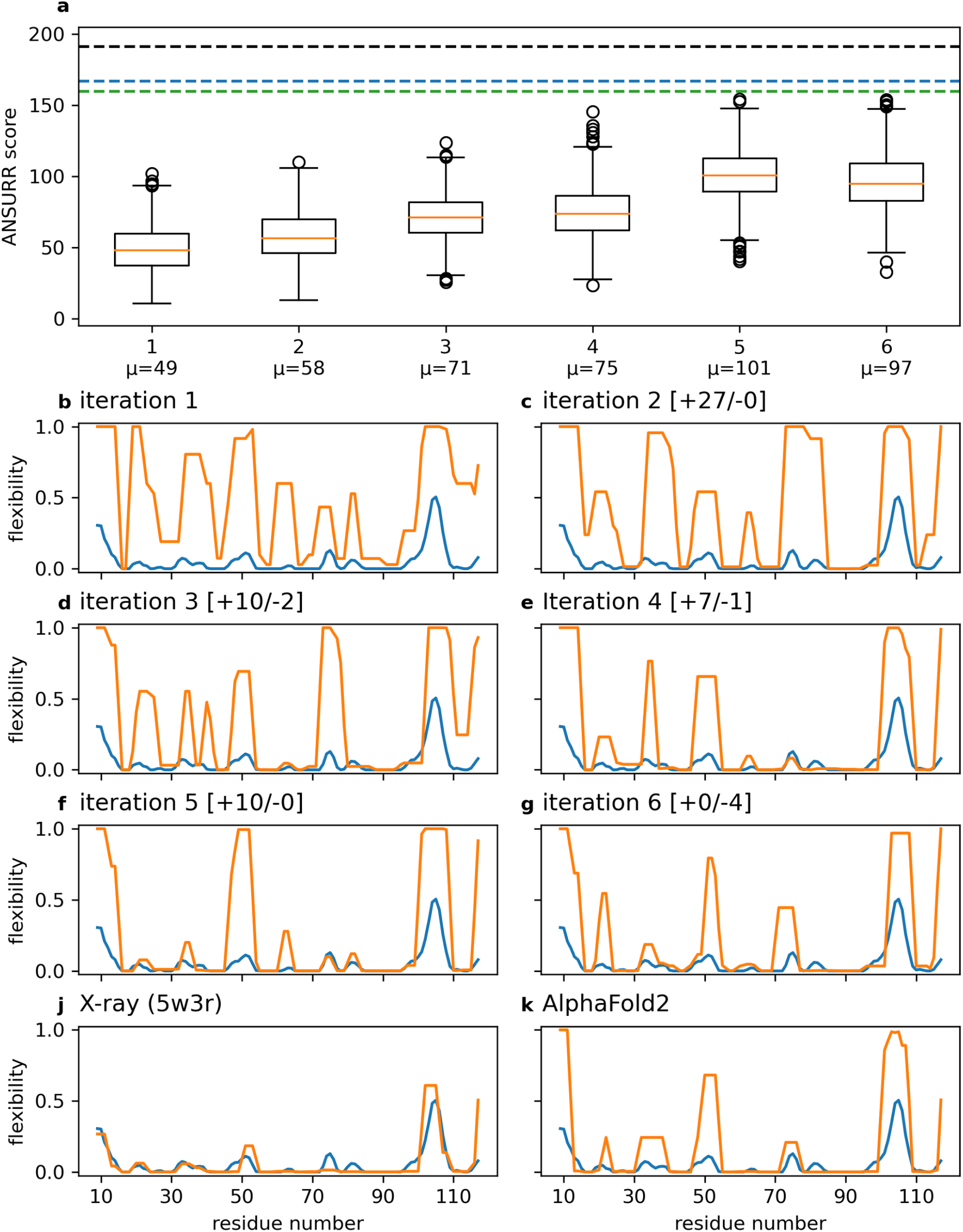
a) Box plots showing the distribution of ANSURR scores for each iteration of CYANA calculations (1200 models in total per iteration). ANSURR scores steadily improve as the set of hydrogen bond restraints is refined, suggesting that inclusion of hydrogen bond restraints does improve structure accuracy. Mean ANSURR scores for each iteration are indicated below the plot. The black dotted line is the score for the X-ray structure 5w3r; the blue dotted line is the mean ANSURR score for 5 models predicted by AlphaFold2 and the green line is 5 models predicted by RosettaFold. b-g) ANSURR comparison of flexibility according to chemical shifts (blue) and flexibility computed for the best scoring model from each iteration (orange). The number of hydrogen bond restraints added and removed between each iteration is indicated in square brackets. j) ANSURR output for the X-ray structure of SH2B1 (PDB ID: 5w3r). k) ANSURR output for the best scoring AlphaFold2 model.

Figure 4b-g depicts the ANSURR output for the best scoring model from each iteration. Fig 4j and 4k show ANSURR output for the X-ray structure (PDB 5w3r) and the best scoring AlphaFold2 prediction, respectively. The crystal structure has a very good ANSURR score, but appears to be more rigid than the “true” solution structure around residues 70-90, judging by the rigidities shown in Figure 4j. With this exception, it would appear to be an excellent guide to the hydrogen bonds present in solution. Indeed, in Figure S4 we present an analysis of the HBRs at each iteration compared to those in the crystal structure, and show that they become progressively closer to those in the crystal structure. We note that the amino acid sequence of the proteins used for NMR and crystallography (mouse and human, respectively) are slightly different (Figure S5), with a total of 5 differences (including the C-terminal residue). It is of interest that the region for which NMR does worst is residues 46-55, which is too flexible in the NMR structures but appears more correctly rigid in the X-ray structure. In the crystal, there is a bound phosphate here from the buffer. This is also where the phosphotyrosine binds. The residues in contact with the phosphate in the crystal structure (residues 47-49) appear to be too rigid, suggesting that the phosphate is more tightly bound in the crystal than in solution, or perhaps that the flexibility according to chemical shifts represents an average of a bound state and unbound state. We have compared backbone chemical shifts for protein in 50 mM Tris buffer and in 50 mM phosphate buffer, pH 6 (Figure S6). It is striking that almost all of the large chemical shift changes surround the location of the phosphate ion in the crystal structure, implying that phosphate binds here in 50 mM phosphate buffer. This may serve to rigidify the region, and explain why the NMR structure determined for protein, which was determined in Tris buffer, has poor rigidity in this region.

### Validation of iterations against common metrics of structure quality

The previous section showed that inclusion of increasing numbers of HBRs generated according to the protocol shown in Figure 2 leads to a steady improvement in ANSURR score: in other words, a more accurate structure. Here, we report on other commonly used measures of quality, to characterise the structural improvement in more detail. The results are shown in Figure 5. The magnitude of NOE violations increases gradually as HBRs are included, although it remains very low throughout (Figure 5a). The calculation of an NMR structure is a balance between different forces pulling in different directions: for example, van der Waals violations push atoms apart, while NOE violations pull them together. For this reason, it is generally found that addition of a new type of restraint leads to a slight increase in violations of the existing restraints (see for example ^38^). We therefore do not see this result as surprising. In a similar way, the magnitude of dihedral angle violations has a small but not significant increase (Figure 5b), as do van der Waals violations (data not shown). The RMSD of backbone atoms to the crystal structure improves on initial addition of HBRs, up to iteration 4 after which there is little change (Figure 5c). This can be understood by the argument that HBRs are excellent for improving local geometry, but make rather little difference to the positions of backbone atoms. It is widely agreed that the template modeling score (TM-score) is a more accurate measure of global similarity than RMSD,^39^ and indeed, we see a more clear trend for TM-score than for RMSD (Figure 5d). All iterations have a TM larger than 0.5, indicating a correct fold. TM-score slightly increases up until iteration 4. Residual dipolar couplings (RDCs) can be used either to validate the quality of NMR structures or as restraints to improve NMR structures, but not both at the same time.^40^ Here we measured a set of ^1^*D*_NH_ as a simple validation measure. Figure 5e shows the RDC *Q* factor,^41^ which improves up to iteration 4. The chemical shifts of amide proteins should be a sensitive measure of structural accuracy, because they are strongly affected by hydrogen bonding.^42^ We therefore calculated the expected amide proton shifts using SHIFTX2^28^ and compared them to the experimental shifts (Figure 5f), which improve steadily as HBRs are added. The distribution of Ramachandran angles also improves slightly up to iteration 4 (Figure 5g). Finally, we have calculated the backbone ensemble RMSD (Figure 5h). This is a measure of precision rather than accuracy, but our survey of PDB structures indicated that it does correlate with accuracy, presumably because adding more (correct) restraints serves to improve both accuracy and precision at the same time;^2^ and indeed, RMSD decreases steadily as HBRs are added. Thus in summary, the gradual addition of 47 HBRs results in an improvement in structural accuracy, as measured by a wide range of parameters. It is worth noting that these metrics clearly show that there is overall no increased penalty to the structure quality from their inclusion: in other words, the metrics shown here imply that the HBRs are correct and do not conflict with the other structural restraints. The structural accuracy clearly improves until iteration 4. Iteration 5 results in an improvement on some measures but a deterioration in others, while iteration 6 (in which some HBRs were removed for lack of good evidence) represents a deterioration in almost all measures, the exception being the precision of the ensemble (Figure 5h).

**Figure 5.**
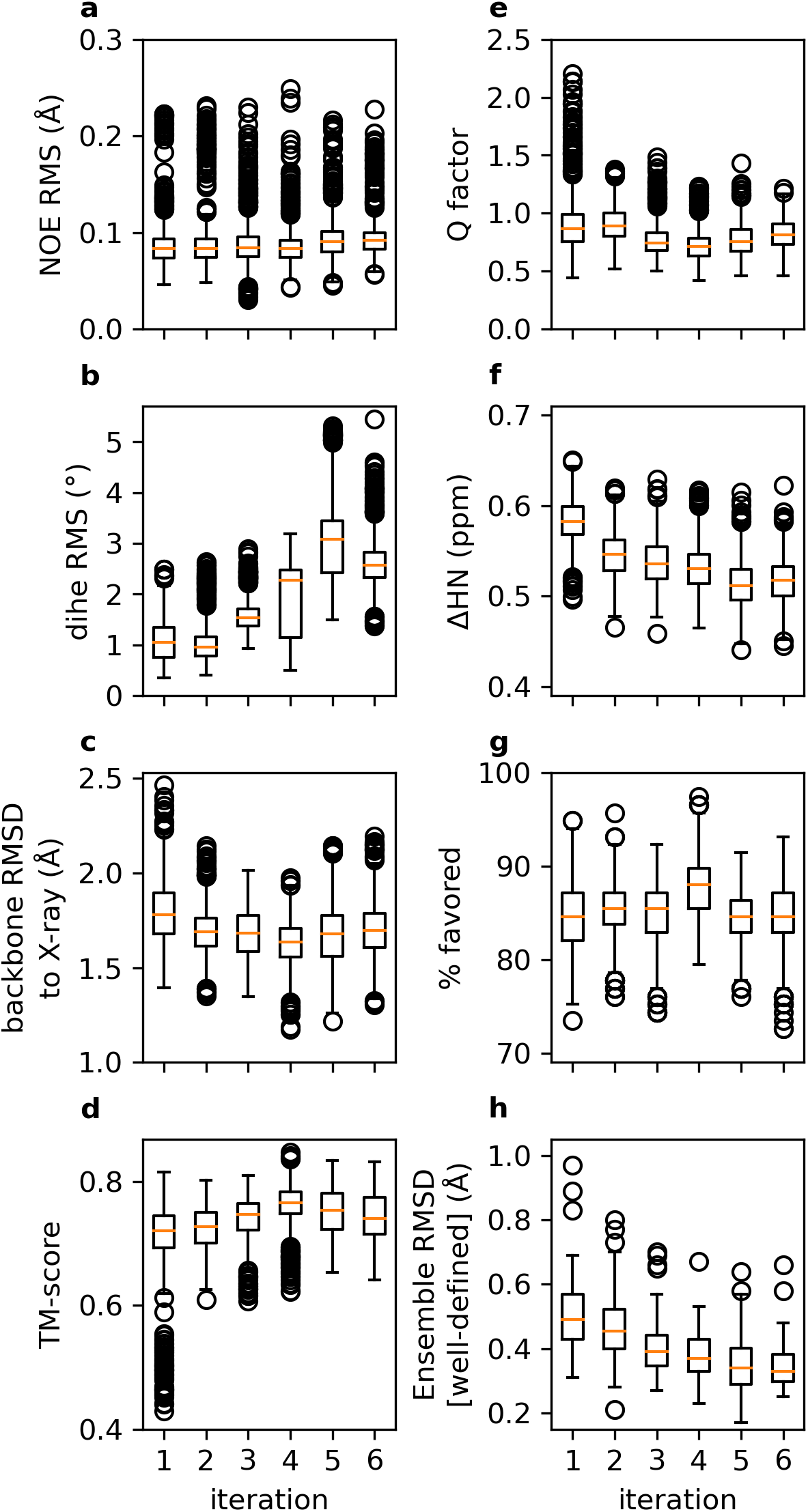
Change in structure metrics during the iterative addition of HBRs. In each panel, the orange bar is the median; box boundaries denote first and third quartiles, and circles denote outliers. Data are shown for each model in the ensemble. (a) Root mean square distance violation. (b) Root mean square dihedral angle violation. (c) Average backbone RMSD to the crystal structure 5w3r. (d) Average template modelling score between NMR and crystal structures, where a value of 1 indicates a perfect match. (e) Residual dipolar coupling *Q* factor. (f) Mean difference between measured amide proton shifts and shifts calculated using SHIFTX2. (g) Percentage of amino acids in the favoured area of the Ramachandran plot. (h) Average backbone RMSD to the mean structure (the ensemble precision).

### Refinement in explicit solvent

It is well established that refinement in solvent is required in order to produce good quality structures.^1, 43^ The 1200 structures produced in each iteration above were therefore refined in water using the program CNS.^20^ There is a reasonable correlation between the energies before and after refinement in explicit solvent (Figure S7). The distance, dihedral angle and hydrogen bond restraints used above were applied during the refinement. Interestingly, if the hydrogen bond restraints were removed at this stage (on the grounds that the structures should now be able to maintain hydrogen bonding geometry because of the force field), hydrogen bonds were lost and the structure quality got worse. This may be an indication that the hydrogen bond potential in CNS is not ideal for this specific purpose: we note a different potential recently introduced into Xplor-NIH that may handle this problem better.^38^ It is encouraging to see that the ANSURR score following refinement in explicit solvent improves (Figure 6, compare to the equivalent results in Figure 4), to the extent that the best scoring structures in the final iterations are competitive with the X-ray structure and AI predictions and could therefore be considered to be of good quality.

**Figure 6.**
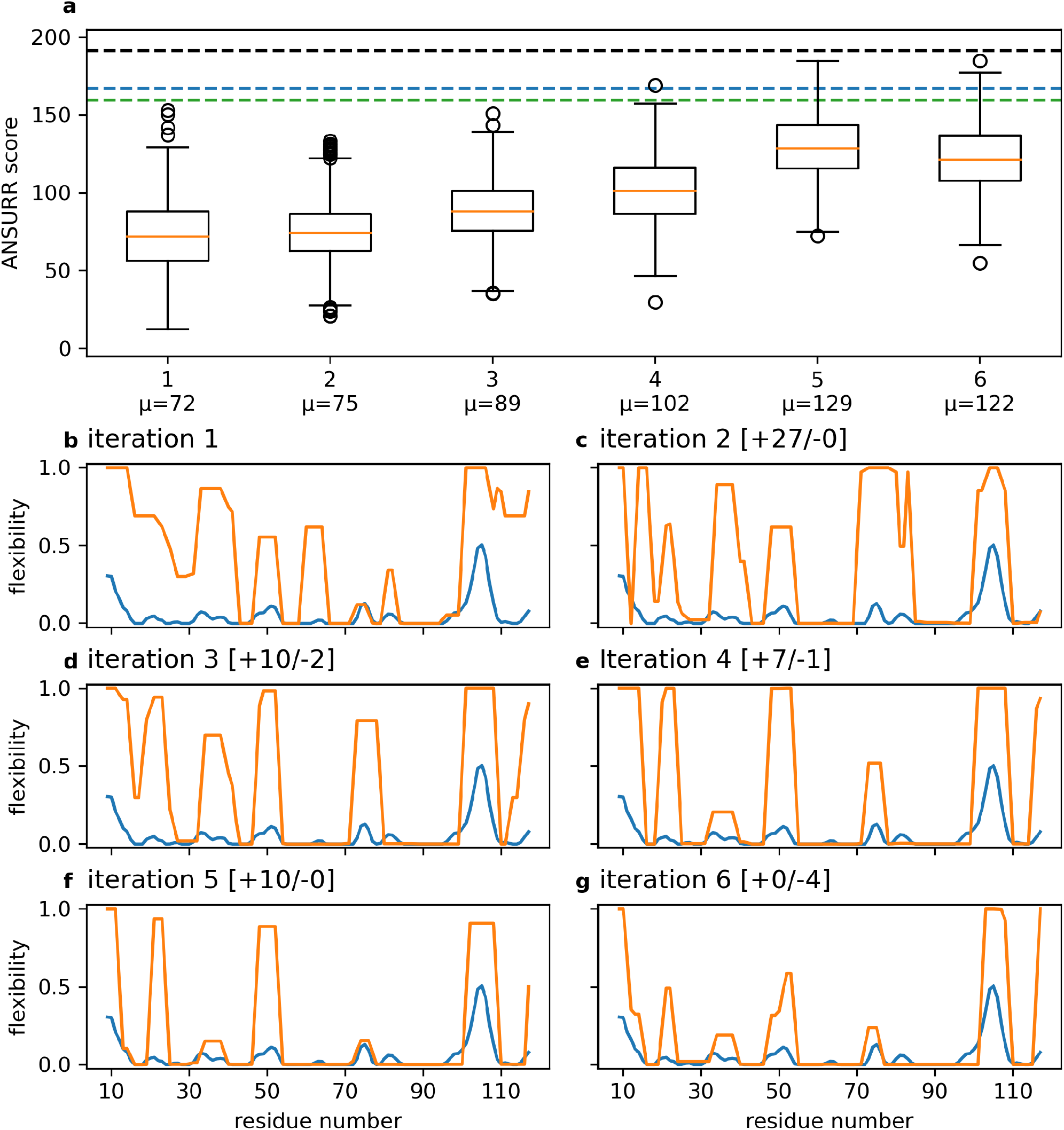
a) Box plots showing the distribution of ANSURR scores for each HBR iteration, following CNS refinement in explicit solvent. These results should be compared to those shown in Figure 4, which show the equivalent results before refinement. The black dotted line is the score for the X-ray structure 5w3r; the blue dotted line is the mean ANSURR score for 5 models predicted by AlphaFold2 and the green line is 5 models predicted by RosettaFold. b-j) Comparison of flexibility according to chemical shifts (blue) and flexibility computed for the best scoring model from each iteration (orange). The number of hydrogen bond restraints added and removed between each iteration is indicated in square brackets.

It is always difficult to know “when to stop” in an NMR structure calculation: improvements and further iterations are always possible. The ANSURR score is a ranked index, compared to all NMR structures in the PDB that we have reliable data for (eg at least 75% completeness of backbone chemical shifts and at least 20 residues long, which comes to 4742 ensembles).^2^ An ANSURR score of 100 is therefore by definition the PDB median; an ANSURR score of greater than 100 is “better than average” and is a good initial goal. We have produced an ensemble with an average ANSURR score of 117, which is well above 100, and with the best members at least as good as AlphaFold2, which we judge to be acceptable.

Despite this, the structure calculated using the final set of 47 HBRs is clearly not “correct” in that the flexibility of residues 46-55 do not match well to the flexibility according to the backbone shifts (Figure 6g). As discussed above, this is probably a binding site for phosphate. It may be that addition of further restraints to a bound phosphate may improve the rigidity here, but this is not justified by the experimental NMR data.

Residues 99-112 are clearly moderately dynamic in solution. The rigidity of the crystal structure matches well to the experimental RCI data (Figure 4j), while unusually,^3^ AlphaFold2 predicts this region to be more flexible than indicated by RCI (Figure 4k). HBRs have produced an overall rigidity in the NMR ensemble that matches the data reasonably well (Figure 6g). However, a comparison of the NMR-derived HBRs with hydrogen bonds in the crystal structure (Figure S8) shows that there are differences in hydrogen bonding in this region. This may indicate some structural heterogeneity in this loop.

The refined structures have been analysed in more detail (Figure 7). The NOE violations in the refined structures (Figure 7a) change very little as HBRs are introduced, suggesting that the HBRs are consistent with the NOE restraints. Calculated amide proton shifts are slightly better in the refined structures (Figure 7f) than the unrefined structures (Figure 5f), as one might expect. The RDC *Q* factors (Figure 7e) improve by about 20% at all iterations. Similarly, Ramachandran distributions are better and more constant in the refined structures (Figure 7g), and RMSDs are rather smaller (Figure 7h). Overall, these statistics show that refinement in water consistently improves the structures at all stages. Taking all measures together, the results imply that adding HBRs in this gradual and validated manner represents a genuine and useful improvement in accuracy.

**Figure 7.**
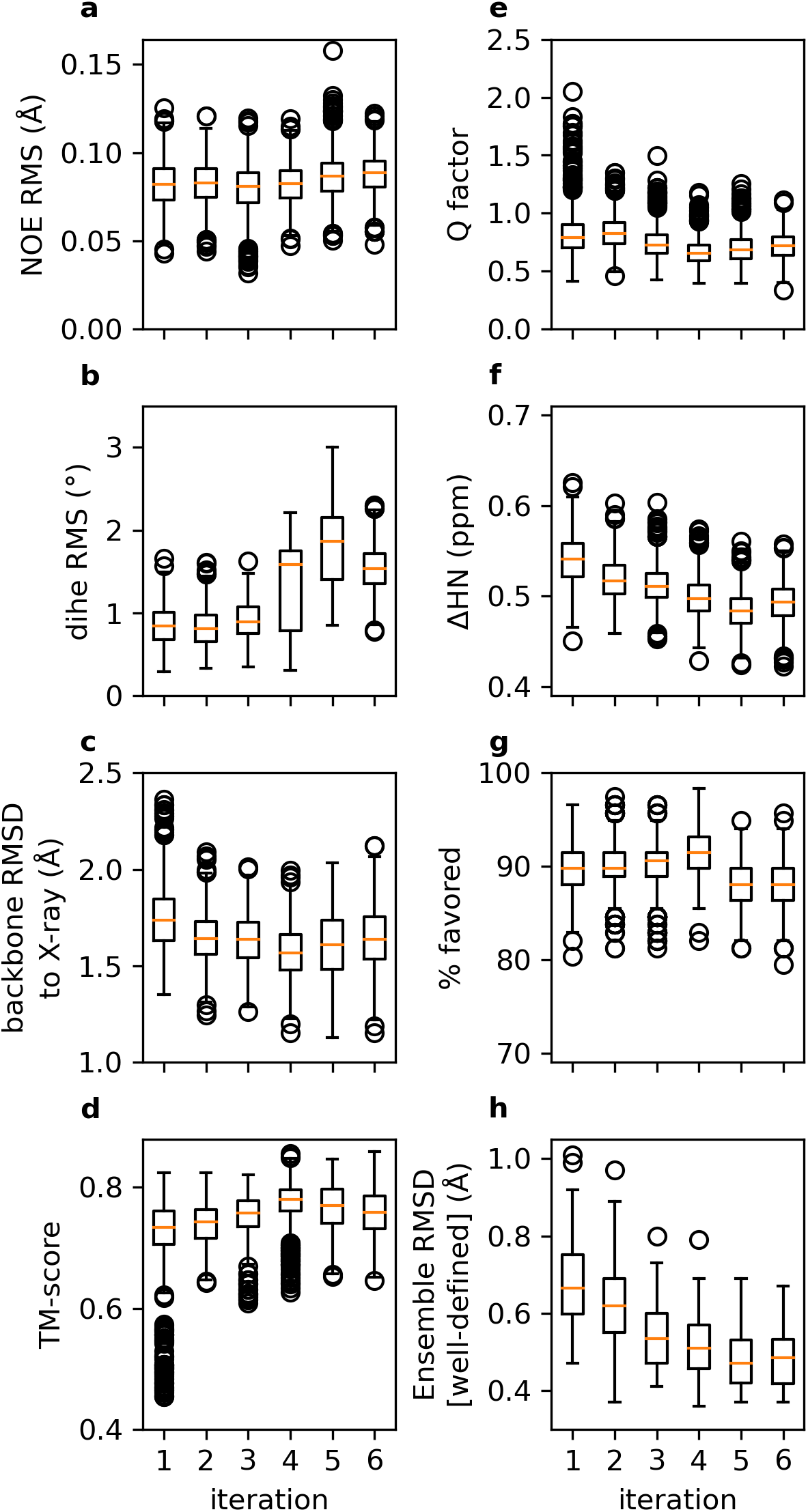
Change in structure metrics of structures refined in water during the iterative addition of HBRs. Compare to Figure 5. (a) Root mean square distance violation. (b) Root mean square dihedral angle violation. (c) Average backbone RMSD to the crystal structure 5w3r. (d) Average template modelling score between NMR and crystal structures. (e) Residual dipolar coupling *Q* factor. (f) Mean difference between measured amide proton shifts and shifts calculated using SHIFTX2. (g) Percentage of amino acids in the favoured area of the Ramachandran plot. (h) Average backbone RMSD to the mean structure (the ensemble precision).

### A consensus NMR ensemble

Each of the 60 CYANA calculations carried out in the final iteration produced a different set of NOE restraints. We investigated several ways to use these to produce a consensus ensemble, and our preferred solution is based on the consensus bundle procedure described by Buchner and Güntert.^36^ In this method, the different sets of NOE restraints are combined to obtain a (smaller) consensus set of NOEs, which are then used to calculate a consensus ensemble. A big advantage of this method is that the RMSD spread (that is, the precision) of this ensemble is <u>larger</u> than is obtained from each of the individual calculations, and is a much more realistic number: in particular, the precision is expected to be of comparable magnitude to the accuracy, as obtained for example by comparing to a crystal structure.

Our consensus NOE restraint set comprised restraints that appeared in at least 36 (60%) of the 60 restraint sets from the final iteration. We then used CYANA to produce 1000 models using only the consensus NOE restraints and TALOS-N dihedral restraints. Finally, we refined the 100 models with the best CYANA target function values in explicit solvent using CNS, adding in the 47 pairs of hydrogen bond restraints from the final iteration, and selected the 20 models with the best NOE energy as our final NMR ensemble. The delayed addition of HBRs reduces the risk of some residues becoming trapped with incorrect torsion angles as CYANA tries to satisfy HBRs at the same time as NOE restraints. This ensemble is shown in Figure 8a. It has been submitted to PDB as ID 8atk. Figure 8b shows the ANSURR scores for these structures, showing a reasonably wide spread of ANSURR scores. We note that the distribution of ANSURR scores for the consensus ensemble is very similar to the ANSURR scores of the individual restraint sets of iteration 6, implying that the use of the consensus restraints has not led to any significant loss in accuracy. We also note that the lowest NOE energy structure (which by default would be model #1 and thus the representative structure of the ensemble) is one of the worst ANSURR scores in the ensemble, while the best ANSURR score is model 18, which is one of the highest NOE energies (although it is worth noting that all NOE energies are very small (Figure 7a) and therefore do not provide much discrimination between structures). The PDB file therefore has model 18 as the first (and by default representative) model in the ensemble.

**Figure 8.**
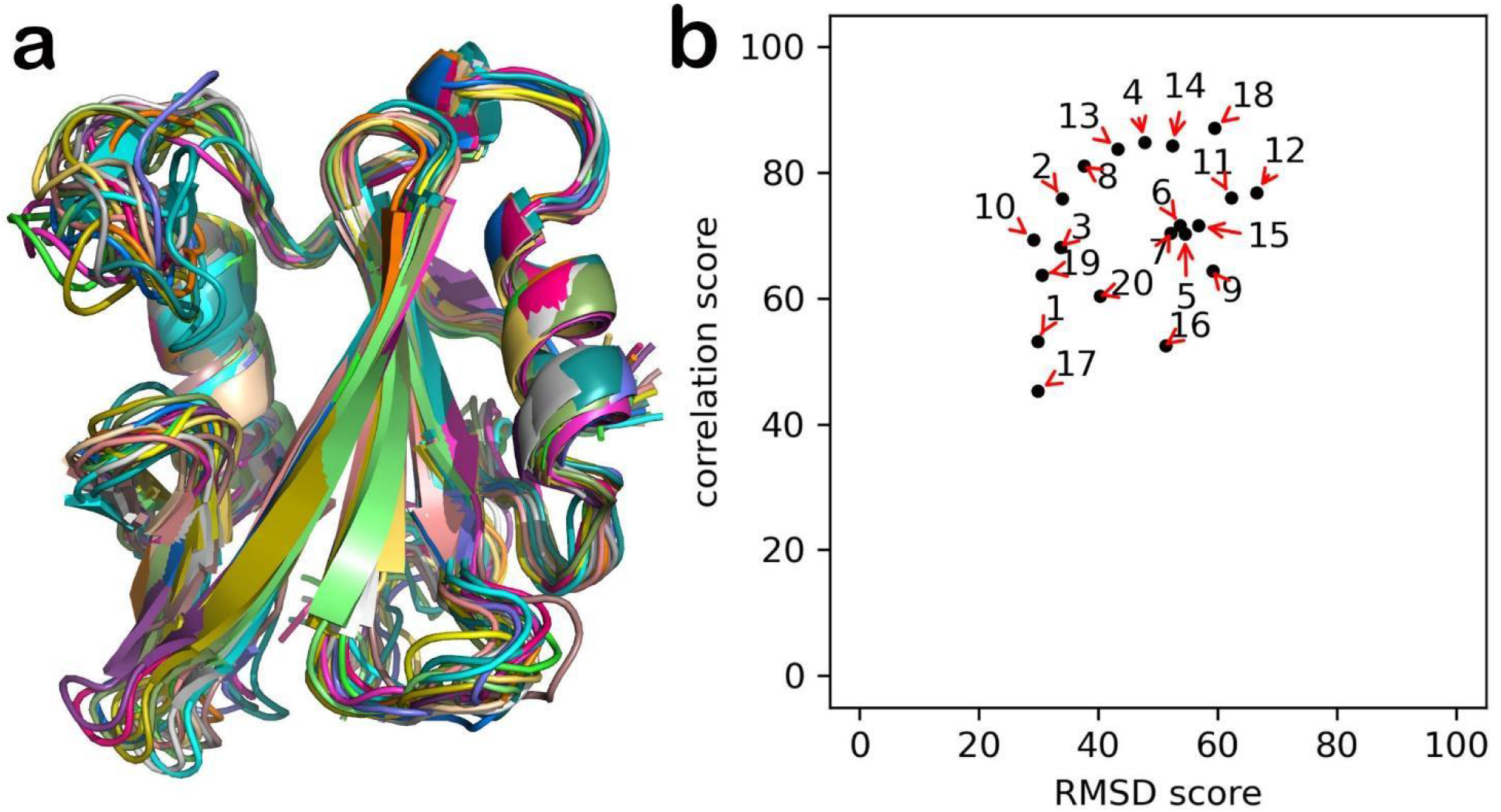
The ensemble formed by selecting the 20 lowest NOE energy models from the final solvent-minimised consensus ensemble. (a) Superposition of the cartoon plot of the structures. In the view shown here, the phosphate binding loop is in the bottom centre, while the dynamic loop 99-112 is top left. The typical SH2 binding site for phosphotyrosine peptides has a pY binding pocket above the phosphate loop, and a second hydrophobic pocket on the left side of the long β-strand at the front. (b) ANSURR scores for these 20 structures. Numbering is in order of NOE energy, with 1 being the lowest.

Table 1 presents a range of statistics for the ensemble. In common with other NMR structure determinations, the Table presents a range of validation measures to enable users to judge the quality of the ensemble. These measures serve different functions: some provide indications of the geometrical qualities of the structures, some provide indications of the accuracy of the structures, and some do both. The input restraints are fairly conventional: no RDCs, and no unusual restraints, but a large number of carefully validated hydrogen bond restraints, which has been the focus of this work. The number and range of restraints encourage us to expect a well-defined but not outstandingly accurate structure: PSVS^44^ calculates 2.9 restricting long-range distance restraints per restrained residue, which is about average. The deviations from idealised geometry provide some reassurance that the structure calculation and energy minimisation process has worked and that the structures are geometrically reasonable, in that there is no evidence of undue strain on the structures; however, they provide no evidence on the accuracy of the structures. The list of restraint violations is a standard requirement and is unremarkable. Because CYANA discards NOE restraints that are repeatedly violated, such a list also provides little guidance on the accuracy of the structure.

**Table 1.**
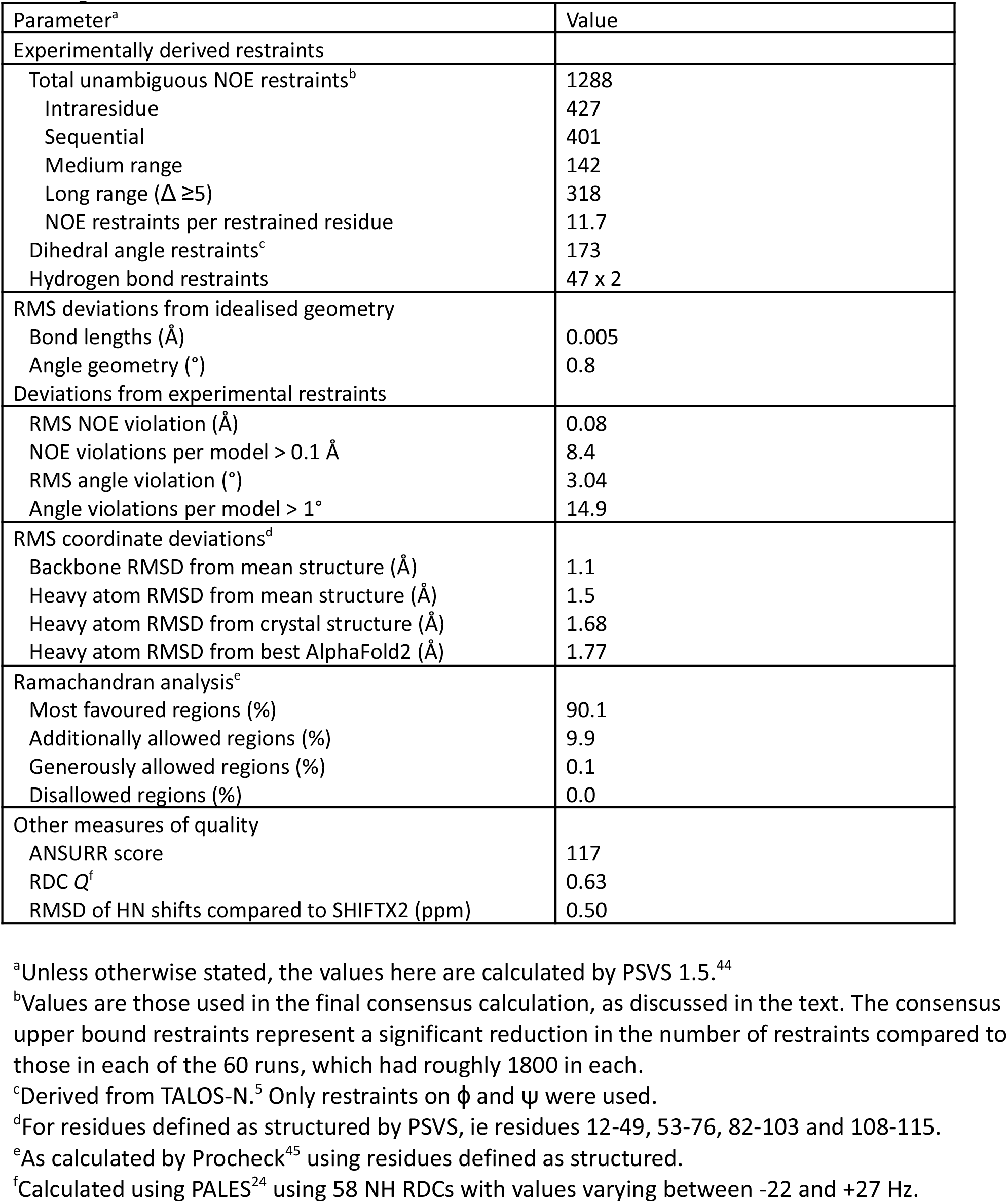
Structural statistics for the SH2 structure ensemble shown in Figure 8. Values are averages over the final ensemble of 20 structures.

The remaining data in Table 1 provide a clearer estimate of the accuracy of the structure. The RMS deviations from the mean structure are explicitly a measure of precision, not of accuracy. However as seen here and elsewhere,^2, 46^ RMSD does correlate with measures of structure accuracy, presumably because an increased number of restraints tends to produce improvements in both accuracy and precision. These RMSD values are rather large for a good structure: however, it is important to note that the consensus calculation adopted here is deliberately aimed at producing as wide a spread of structures as possible, and should ideally give a precision that is close to the true accuracy.^36^ It is widely assumed that the crystal structure is the closest one can get to the “correct” solution structure. Our work on ANSURR^1-2^ has shown that the crystal structure is generally an accurate representation of the solution structure, though it tends to be too rigid. ANSURR reports that in general the crystal structure is considerably more accurate than the NMR ensemble; and a survey from the Montelione lab^47^ showed that the RMSD precision is usually tighter than the average distance between the NMR ensemble and the crystal structure (the “accuracy”), by a factor of 2.7 ± 1.3. We thus expect the RMS distance to the crystal structure to be small, and ideally of the same magnitude as the consensus ensemble RMSD. Given the remarkable success of AlphaFold2 in predicting structures accurately^3, 48-49^, we have also included an RMSD to AlphaFold, with similar expectations. The observation that these RMSD values are small, and of similar magnitude to the RMSD precision, is a good indication of the accuracy of the structures, and suggests that the precision obtained from the consensus ensemble is a better guide to the true accuracy, as claimed.^36^

In principle, the Ramachandran distribution tells nothing about the accuracy of the structure, but merely reports on whether it has good geometry. However, our experience with ANSURR^1-2^ demonstrates that the Ramachandran score is additionally a remarkably good guide to accuracy. ANSURR itself reports that the ensemble is good but not excellent, with some members of the ensemble having high accuracy (Figure 8b). Our experience from this work and earlier studies^1-2^ thus encourages us to see Ramachandran and ANSURR scores (together with the RMS to the crystal structure) as the best measures for validating the accuracy of the NMR ensemble. In support of this assertion, we have compared ANSURR scores to energies for the members of the final iteration, shown in Figure 9. The data show a highly significant correlation between the two, encouraging us that ANSURR provides a useful validation of accuracy. Table 1 also includes a comparison of experimental and back-calculated amide proton shifts. This measure improved consistently during the iterations and could prove to be a simple and useful guide to accuracy

**Figure 9.**
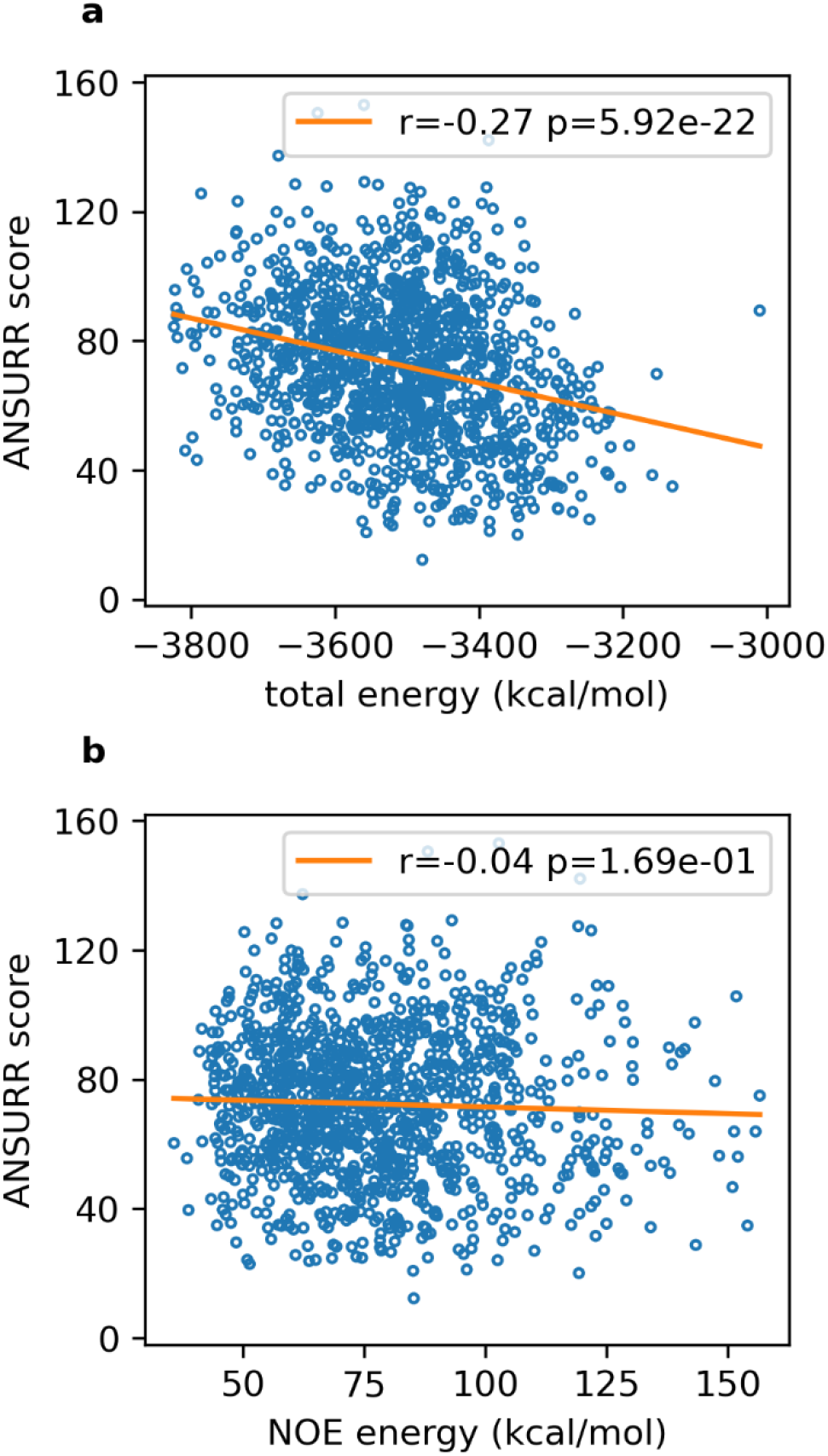
Correlation between ANSURR score and energy, for the 1200 structures calculated in the final iteration and refined in explicit solvent. The orange line is the line of best fit, and the figures give the Pearson correlation coefficient and its two-tailed *p* value. (a) Total energy (b) NOE energy.

## Discussion

Protein structure determination by X-ray crystallography uses the position and intensity of diffraction data as its input. Because there is a direct relationship between structure and diffraction pattern, the same data can also be used to check the accuracy of the resulting structure, via the *R* and *R*_free_ factors. NMR input data is much smaller in number, less precise and more diverse. There is also no simple relationship between the input data and output structure. The consequence is that structure calculation by NMR relies on generating an appropriate combination of different structural restraints, including a heavy reliance on knowledge-based restraints, where part of the challenge is to introduce and weight the different types of restraints appropriately. NMR has always had a problem with structure validation.^50^ Here, we have taken the approach that some types of data (NOEs, dihedral restraints derived from backbone shifts, and hydrogen bonds inferred using a combination of temperature coefficients and slow exchange) are used to generate restraints and/or structure selection criteria; while others (ANSURR scores based on rigidity, Ramachandran distribution, residual dipolar couplings [RDCs], back-calculated amide shifts, and [where appropriate] comparison to crystal or AlphaFold structure) are used to measure accuracy and <u>not</u> to produce restraints. If any of this second group is used to drive structure calculation or selection, then clearly it can no longer be used as a measure of accuracy. This classification is not unique: there are for example many structure calculations that have successfully used RDCs as restraints, in which case they should not be used to measure accuracy.

In this paper, we have proposed an iterative protocol for identifying hydrogen bond restraints based on experimental NMR data. The HBRs identified in this way were compatible with the NOE and dihedral restraints and significantly improved the overall accuracy of the structures. The resulting ensemble had a lower energy, better ANSURR score, better Ramachandran distribution, better similarity to the crystal structure, better RDC *Q* factor, and lower RMSD than ensembles calculated with no or fewer HBRs. We suggest that this protocol could be used as the basis for a robust, transparent, and automatable system for calculating better NMR structure ensembles. We note several key features of the protocol: HBRs were introduced gradually and iteratively, with a final step where possible incompatible HBRs were removed; the initial calculations used only NOEs and dihedral restraints; and multiple ensembles were calculated at each stage, to obtain better distinction between correct and incorrect HBRs. The final calculation used consensus distance restraints in order to sample conformational space better and obtain a more realistic and useful RMSD.^36^

Only backbone-backbone HBRs were applied here. One could easily imagine adding sidechain restraints, although in most cases the experimental evidence required might be harder to obtain. For SH2, there are two pairs of sidechain hydrogen bond restraints for which we have clear evidence. (a) The sidechain amides of Asn59 are very slowly exchanging. In our structures, they are buried inside the structure and both form well-defined hydrogen bonds to the backbone carbonyls of V31 and G35, even without restraints being added. Thus the restraints could be added, but would actually not make any difference, because hydrogen bonding is already present due to the local density of restraints. (b) It is well established that α-helices often have caps at both ends, but particularly at the N-terminal end where sidechains of Ser, Thr or Asn form hydrogen bonds to (*i*+4) residues within the helix.^51^ Both of the SH2 helices have serines as the N-capping residue, and slowly exchanging amides at the relevant positions. We have recalculated ensembles with these N-cap restraints in place. Structures are well formed and of low energy, but the added restraints make very little difference to the overall energy or accuracy.

Many of the validation measures used show an improvement in accuracy with addition of extra HBRs (Figures 5 and 7). In the absence of a “true” structure for comparison, the best measures of accuracy across the entire refinement process appear to be ANSURR score and Ramachandran distribution, with RDCs some way behind. Ramachandran is not an ideal measure because it measures how good a protein structure it is, rather than how well it matches experimental data, and therefore is liable to give misleading results if the calculation is heavily biased towards known “good” structures. RDCs are useful for validation, though not if they are already being used as restraints. Compared to ANSURR, they lack power to differentiate between poorer quality structures ^48^. We therefore propose the ANSURR score as a useful, simple and rapid measure of accuracy. In particular, we noted above that by definition an ANSURR score of 100 is an “average” accuracy. Thus, any structure ensemble that achieves an ANSURR score of greater than 100 is “better than average” and is at least acceptable. If all new NMR structures in the PDB had an ensemble ANSURR score greater than 100, this would lead to a gradual overall improvement in the accuracy of NMR structures in the PDB.

During the development of the iteration protocol, we deliberately did not compare our hydrogen bonds to those observed in the crystal structure. On making such a comparison (Figure S8), there is in general a remarkably good agreement between the hydrogen bonds in the crystal and those identified by the procedures detailed here, especially for the regular secondary structure elements.

One might imagine that incorporation of explicit HBRs is not necessary, in that if a structure is almost correct, then refinement in explicit solvent should drive the formation of the correct hydrogen bonds. However, this does not in general seem to be true. There is some improvement in hydrogen bonding arising from structure refinement,^52^ but it seems that the geometry has to be very close to correct in order for the forcefield to produce a hydrogen bond. We note that Schwieters et al^38^ have introduced a new potential into Xplor-NIH that may improve the situation; and that AI-based methods such as AlphaFold have good success in predicting hydrogen bonds. Nonetheless, it is currently important to maximise the number of correct HBRs in NMR structure calculation. We hope that the proposals outlined here will help.

## Conclusions

The future of biological NMR as a structural tool in the post-AlphaFold era lies in its ability to detect and characterize structural and dynamic heterogeneity in solution. To carry out that task effectively it is essential that the structural details from NMR should be as accurate as possible. Here we have shown that hydrogen bond restraints can be added in a transparent and justified way, and that these significantly improve the accuracy of the structure. We have also shown that ANSURR is a useful tool for assessing accuracy, and can be used to adjudge when the structure determination process is ready to complete.

## Associated content

### Supporting information

The supporting information is available free of charge at xxxx.

Three Tables (list of matching crystal and NMR structures; NMR experimental details; hydrogen bond restraints in each iteration) and 8 figures.

### Accession codes

The coordinates of the final set of 20 structures have been deposited in the Protein Data Bank under code 8atk, and NMR chemical shifts have been deposited in BioMagResBank under code 51342.

## Acknowledgements

We thank the Biotechnology and Biological Sciences Research Council for funding to N. J. F. (BB/P020038/1) and for support to upgrade the NMR spectrometer (BB/R000727/1).

