## Supplementary Information for "Improved methodology for protein NMR structure calculation using hydrogen bond restraints and ANSURR validation: the SH2 domain of SH2B1"

### Supplementary Material

**Table S1.** The 215 pairs of NMR and crystal structures used for the comparison in Figure 1. The table lists the Uniprot accession number, and the PDB code, including which chain in the PDB was used for the comparison.

| UP | NMR | XRAY |
|---|---|---|
| P12497 | 2LF4_A | 3H4E_F |
| P01040 | 1CYU_A | 3KSE_D |
| P04271 | 2PRU_A | 3D10_A |
| P04637 | 2MZD_B | 2B3G_B |
| Q9S446 | 1IJA_A | 1T2W_A |
| P00441 | 2AF2_A | 4A7U_A |
| P29160 | 1GO0_A | 2BO1_A |
| P63316 | 2N79_C | 1J1D_A |
| P0DP23 | 2KUG_A | 4DJC_A |
| Q9UKL4 | 2N6A_A | 4DJC_A |
| P61769 | 2XKS_A | 1K5N_B |
| Q00987 | 2LZG_A | 4HBM_A |
| P05112 | 1BCN_A | 2D48_A |
| P00644 | 2KHS_A | 2SNS_A |
| P19909 | 2N9L_A | 3FIL_A |
| P56552 | 2LY5_A | 4HE7_A |
| Q09472 | 2K8F_A | 3IO2_A |
| Q12923 | 3PDZ_A | 3LNY_A |
| P20701 | 1DGQ_A | 1RD4_C |
| P39900 | 2MLS_A | 1JIZ_A |
| O60885 | 2MJV_B | 5U2C_A |
| P98170 | 2ECG_A | 5O6T_A |
| Q15109 | 2L7U_A | 3O3U_N |
| P61956 | 5GHB_A | 5D2M_E |
| P0CG47 | 2KWU_B | 5NVG_A |
| P16333 | 2JS2_A | 5QU4_B |
| Q12809 | 2L0W_A | 4HQA_A |
| Q13148 | 2N4P_A | 5MDI_B |
| Q96T88 | 2LGK_A | 3ASL_A |
| Q9NP97 | 1Z09_A | 2HZ5_A |
| Q9GZQ8 | 2N9X_A | 2ZJD_C |
| P48059 | 2KBX_B | 4HI9_B |
| P06768 | 1B4M_A | 1OPA_A |
| P45481 | 2LQH_A | 4I9O_A |
| P00947 | 1BUQ_A | 1OHP_A |
| P26599 | 1SJR_A | 3ZZY_A |
| P45452 | 1FM1_A | 5B5O_B |
| P38505 | 2M7K_A | 2NXQ_A |
| P41784 | 2KV7_A | 3ZQE_A |
| P02511 | 2N0K_A | 4M5S_A |
| O75369 | 2EE9_A | 5DCP_A |
| O14907 | 2L4T_A | 4NNM_A |
| Q58AD3 | 2KM0_A | 3DSO_A |
| P06241 | 2MRJ_A | 4U17_B |
| O35820 | 2KHZ_A | 4P5D_C |
| P06702 | 5I8N_A | 4GGF_T |
| P14120 | 1CN7_A | 1NMU_B |
| P18887 | 1XNT_A | 3LQC_A |
| Q12824 | 5L7B_A | 5GJK_B |
| Q99986 | 2KUL_A | 5UKF_A |
| P62826 | 2MMC_A | 4HAT_A |
| Q03526 | 2K7A_B | 3S9K_A |
| Q92793 | 2LXS_A | 5SVH_A |
| P23528 | 1Q8X_A | 4BEX_1 |
| P06766 | 1DK2_A | 1BPD_A |
| P09038 | 1BLD_A | 1IIL_A |
| Q8KC80 | 2KO1_A | 3IBW_A |
| P01563 | 2LMS_A | 4YPG_D |
| P0AA04 | 2LRK_D | 3CCD_A |
| O15151 | 2N14_A | 3FDO_A |
| P32851 | 1BR0_A | 1EZ3_C |
| P47992 | 2N54_A | 2NYZ_E |
| Q13526 | 2RUR_A | 3I6C_B |
| P99999 | 2N9I_A | 5TY3_A |
| P26447 | 2MRD_A | 4CFR_B |
| P0C077 | 2KC8_A | 4FXI_B |
| P56945 | 5O2M_A | 1WYX_B |
| P29375 | 2KGG_A | 3GL6_A |
| P01139 | 5LSD_A | 1SGF_Y |
| P0C6U8 | 2LIZ_A | 3VB5_B |
| P31948 | 2NC9_A | 1ELR_A |
| Q13158 | 1E41_A | 3EZQ_D |
| O95931 | 2K1B_A | 4MN3_A |
| Q14527 | 5K5F_A | 4XZG_C |
| O85142 | 2M30_A | 1R1U_D |
| Q9H3D4 | 1RG6_A | 2Y9U_A |
| P63165 | 5GHD_A | 1Y8R_C |
| O95786 | 2LWE_A | 4P4H_F |
| P03495 | 1NS1_A | 1AIL_A |
| Q9Y294 | 1TEY_A | 2IO5_A |
| O75475 | 2N3A_B | 2B4J_C |

Q03164 2KYU_A 3LQH_A
P08603 2JGW_A 2UWN_A
P04925 2L39_A 4MA7_A
P9WI81 2KUD_A 5E10_A
P01344 2L29_B 3KR3_D
P22217 2N5A_A 3F3Q_A
P12689 5VX7_A 5UMV_A
P55059 2DJK_A 5CRW_A
P11940 2RQG_B 3KUT_A
Q92784 2KWN_A 5SZC_A
Q9UER7 2KZU_A 5GRQ_A
P15692 1KAT_V 4ZFF_D
P27577 1R36_A 1MD0_A
P04156 1QM1_A 3HAK_A
P80511 2M9G_A 2WCB_A
Q00459 1G11_A 3GE3_E
Q9JIL4 2EDZ_A 3NGH_B
P03052 2N5G_A 5CKT_A
P0AFU8 1I18_A 1I8D_A
Q9GTP0 1EWW_A 1L0S_D
Q92859 1X5J_A 4UI2_A
P08839 3EZB_A 2HWG_B
Q12285 4ASW_C 4A20_A
Q9UKV5 2LVO_C 4G3O_A
Q14160 1UJU_A 4WYU_B
O95166 1KLV_A 1GNU_A
P07602 1SN6_A 2GTG_A
P68390 1M8B_A 1PPF_I
Q99814 5KIZ_A 3F1P_A
Q251Q8 2KYI_A 3IPF_A
Q92541 2DB9_A 4L1P_B
Q5VST9 2MWC_A 4RSV_A
Q5W1E8 1XSX_A 1R7J_A
Q99584 1YUU_A 2H2K_B
Q9Y530 2L8R_A 4J5Q_A
Q05H60 2LPI_A 3LR2_B
O28769 2L7H_A 2Y20_B
Q8IUC6 2M63_A 4BSX_B
P62495 2HST_A 1DT9_A
P50583 1XSB_A 4ICK_A
O34816 3ZQD_A 4A1K_A
O14745 2JXO_A 4Q3H_A
Q84B82 2KMT_A 3TCJ_A
Q79PF6 1M2F_A 5C5E_B
Q16674 1HJD_A 5IXB_A
P78310 1RSF_A 1KAC_B
Q99816 1M4P_A 4YC1_A
P21457 1JSA_A 1OMR_A
P09237 2DDY_A 1MMR_A
O66551 2JRL_A 3DZD_B
P36328 5MQX_A 3GQE_A
P14204 2KRF_A 3ULQ_B
O95319 2MY7_A 4TLQ_B
Q14498 2MHN_A 4YUD_A
P26039 2KBB_A 4F7G_B
P22069 2LT5_A 3SNF_A
P02749 1G4F_A 1C1Z_A
P80226 2LFO_A 1TW4_B
P21579 2K45_A 2R83_B
P77173 1F7X_A 1F47_B
P55854 2MP2_A 2IO1_D
P03012 1HX7_A 2RSL_C
P45947 1Z2E_A 1JL3_B
P48061 2KED_A 4UAI_A
P0A744 2GT3_A 1FF3_A
Q96FQ6 2L51_A 3NXA_A
P05434 2AMI_A 3QRX_A
P00257 1L6V_A 1E6E_B
P13479 1E0H_A 2VLQ_A
Q79V62 1Q6B_A 5JWR_F
P43146 2EDD_A 4URT_B
P34174 1LS8_A 2FJY_B
P49960 2GO9_A 2GHP_F
Q96JM7 1WJS_A 3UT1_A
O60260 1IYF_A 5C1Z_B
P03960 1U7Q_A 5MRW_J
P0ACX3 2ASY_A 1WD6_A
Q6R3M4 2KWU_A 3AI4_A
P02696 1MX7_A 1CRB_A
Q8VL32 2GZU_A 2XI8_B
P0ABD8 2BDO_A 1BDO_A
P27540 2HV1_A 3F1P_B
P38182 2KWC_A 3RUI_B
P35225 1IK0_A 3LB6_A
P07854 2FVA_A 2XZ0_D
Q07157 2JWE_A 2RCZ_B
Q16186 2NBV_A 5V1Y_B
Q9BX66 2DL3_A 4LNP_A
O97111 2LC7_A 1RZX_A
O22265 2HUG_A 5E4W_D
P46379 1WX9_A 4DWF_B
P07992 2JPD_A 2A1I_A
Q5PSJ1 1Z7P_A 2E7P_C
P27439 3MSP_A 2BVU_D
P98066 2N40_A 2PF5_D
P00381 2L28_A 3DFR_A
G2EA45 2MPB_A 4OA3_A
Q93IE7 2RVA_A 4ZZ5_B
O35987 1I42_A 1S3S_G
Q12118 2LXB_A 3ZDM_A
Q97ZF4 2A2Y_A 1UDV_A
A0A0H2UPA7 2L3A_A 4G06_B
P37554 2RO5_A 2W1T_A
Q9LUV2 1Q53_A 1Q4R_A
Q9USH8 2LVX_A 4XQM_A
P29355 1KFZ_A 2SEM_B
Q07955 2M7S_A 4C0O_C
Q79V61 5JYV_B 2QKE_B
P0A734 1EV0_A 3R9J_D
P03126 2LJY_A 4XR8_F
O43639 1Z3K_A 2CIA_A
P14949 2GZZ_A 2VOC_A

P25638 2LSV_A 4CGU_A
Q9X0J6 1LKN_A 1O5U_B
P07237 1BJX_A 4JU5_A
P52735 2LNW_A 4ROJ_C
O59169 2M3X_A 4UZR_B
P08622 1BQ0_A 5NRO_B
P0AEC3 1FR0_A 2A0B_A
O82040 2LVJ_A 1K9U_A
Q9BTP7 2M9N_A 4M6W_B
Q86PR3 2KSI_A 1PZ4_A
P49916 1IN1_A 3PC7_A
P02692 2JU8_A 1LFO_A
Q6N882 2LL8_A 3LMO_A
Q8EF26 2JUW_A 2QTI_A
Q7A260 1KTU_A 2O3B_B
P49799 1EZY_A 1AGR_E
O77077 2FQ2_A 3GZM_B
B3GA02 2LIQ_A 4USO_A
P97924 2KR9_A 5O33_B
Q481E4 2JR2_A 2OTA_A
Q0SXH8 1YZA_A 4N4W_A
Q920Q2 2LSG_A 4FJO_A

**Table S2.** Experimental parameters for multidimensional NMR spectra used to assign backbone and sidechain of SH2, and provide NOE restraints for structure calculation. The spectrometer frequency was 600 MHz $^{1}$H. Nuc: nuclei recorded, Sw: spectra width, TD: size of fid, Acq: acquisition time.

| Experiment | Bruker pulse prog | D1 | | | | D2 | | | | D3 | | | |
|---|---|---|---|---|---|---|---|---|---|---|---|---|---|
| | | Nuc | Sw Hz | TD | Acq s | Nuc | Sw Hz | TD | Acq s | Nuc | Sw Hz | TD | Acq s |
| Backbone | | | | | | | | | | | | | |
| N$^{15}$ HSQC | hsqcetfpf3gp | $^{1}H_N$ | 9615.3 | 2048 | 0.1065 | $^{15}N$ | 2189.4 | 256 | 0.0584 | | | | |
| HNCO | hncogp3d.2 | $^{1}H_N$ | 9615.3 | 1024 | 0.0532 | $^{15}N$ | 2189.4 | 40 | 0.0091 | $^{13}C$ | 1811.1 | 100 | 0.0276 |
| HN(CA)CO | hncacogp3d.2 | $^{1}H_N$ | 9615.3 | 1024 | 0.0532 | $^{15}N$ | 2189.4 | 40 | 0.0091 | $^{13}C$ | 1811.1 | 100 | 0.0276 |
| HNCA | hncagp3d.2 | $^{1}H_N$ | 9615.3 | 1024 | 0.0532 | $^{15}N$ | 2189.4 | 40 | 0.0091 | $^{13}C$ | 4527.3 | 100 | 0.0110 |
| HN(CO)CA | hncocagp3d.2 | $^{1}H_N$ | 9615.3 | 1024 | 0.0532 | $^{15}N$ | 2189.4 | 40 | 0.0091 | $^{13}C$ | 4527.3 | 100 | 0.0110 |
| HNCACB | hncacbgp3d.2 | $^{1}H_N$ | 9615.3 | 1024 | 0.0532 | $^{15}N$ | 2189.4 | 40 | 0.0091 | $^{13}C$ | 10563.6 | 160 | 0.0075 |
| CBCA(CO)NH | hncocacbgp3d.2 | $^{1}H_N$ | 9615.3 | 1024 | 0.0532 | $^{15}N$ | 2189.4 | 40 | 0.0091 | $^{13}C$ | 10563.6 | 160 | 0.0075 |
| Sidechain | | | | | | | | | | | | | |
| $^{13}$CHSQC | hsqcctetgpsisp | $^{1}H_N$ | 9615.3 | 1024 | 0.0532 | $^{13}C$ | 12072.7 | 256 | 0.0106 | | | | |
| HBHAcoNH | hbhaconhgpwg3d | $^{1}H_N$ | 9615.3 | 2048 | 0.1065 | $^{15}N$ | 2189.4 | 40 | 0.0091 | $^{1}H$ | 9615.3 | 128 | 0.0066 |
| CcoNH | ccconhgp3d.2 | $^{1}H_N$ | 8196.7 | 2048 | 0.1249 | $^{15}N$ | 2189.4 | 40 | 0.0091 | $^{13}C$ | 12072.7 | 128 | 0.0053 |
| HCcoNH | hccconhgpwg3d2 | $^{1}H_N$ | 9615.3 | 2048 | 0.1065 | $^{15}N$ | 2189.4 | 40 | 0.0091 | $^{1}H$ | 9615.3 | 128 | 0.0066 |
| HCCHTOCSY | hcchdigp3d2 | $^{1}H_C$ | 8196.7 | 2048 | 0.1249 | $^{13}C$ | 12072.7 | 64 | 0.0026 | $^{1}H$ | 8196.7 | 128 | 0.0078 |
| HCCHCOSY | hcchcogp3d | $^{1}H_C$ | 8196.7 | 2048 | 0.1249 | $^{13}C$ | 12072.7 | 64 | 0.0026 | $^{1}H$ | 8196.7 | 128 | 0.0078 |
| Aromatic | | | | | | | | | | | | | |
| $^{13}$C HSQC/TROSY | trosyargpphwg | $^{1}H_C$ | 8196.7 | 2048 | 0.1249 | $^{13}C$ | 5131.3 | 160 | 0.0155 | | | | |
| HBCBCGCDHD | hbcbcgcdhdgp | $^{1}H_{\delta}$ | 9615.3 | 2048 | 0.1065 | $^{13}C_{\beta}$ | 3621.7 | 80 | 0.0110 | | | | |
| HBCBCGCDCE | hbcbcgcdcehegp | $^{1}H$ | 9615.3 | 2048 | 0.1065 | $^{13}C_{\beta}$ | 3621.7 | 80 | 0.0110 | | | | |
| $^{1}$H $^{1}$H NOESY | noesyphpr | $^{1}H_N$ | 7462.6 | 2048 | 0.1372 | $^{1}H$ | 7501.6 | 512 | 0.0341 | | | | |
| Structural (restraint) | | | | | | | | | | | | | |
| $^{15}$N NOESY | noesyhsqcfpf3gpsi3d | $^{1}H_N$ | 9615.3 | 2048 | 0.1065 | $^{15}N$ | 2189.4 | 40 | 0.0091 | $^{1}H$ | 9602.1 | 128 | 0.0066 |
| $^{13}$C NOESY | noesyhsqcetgpsi3d | $^{1}H_C$ | 9615.3 | 2048 | 0.1065 | $^{13}C$ | 12072.7 | 64 | 0.0026 | $^{1}H$ | 9602.1 | 128 | 0.0066 |
| $^{13}$C NOESYaro | noesyhsqcetgpsi3d | $^{1}H_C$ | 9615.3 | 2048 | 0.1065 | $^{13}C$ | 5131.3 | 40 | 0.0038 | $^{1}H$ | 9602.1 | 128 | 0.0066 |

**Table S3.** Hydrogen bond restraints used in the iterative structure calculation. Iteration #1 had no hydrogen bond restraints. To make it easier to identify, at the first iteration where a new hydrogen bond is included as a restraint, it is denoted New.

| Hydrogen bond | | Iteration number | | | | |
|---|---|---|---|---|---|---|
| HN | CO | 2 | 3 | 4 | 5 | 6 |
| G14 | L12 | | | | New | X |
| Y15 | L12 | | | | New | X |
| A27 | S23 | New | X | X | X | X |
| A28 | R24 | | | New | X | X |
| Q29 | L25 | New | X | X | X | X |
| L30 | K26 | New | X | X | X | X |
| V31 | A27 | New | X | X | X | X |
| L32 | A28 | New | X | X | X | X |
| E33 | L30 | New | X | X | X | |
| G35 | L32 | New | X | | | |

| | | | | | | |
|---|---|---|---|---|---|---|
| S38 | G35 | New | x | X | X | X |
| H39 | T36 | | New | X | X | X |
| G40 | N59 | | | | New | X |
| V41 | S38 | New | X | X | X | X |
| F42 | S113 | | New | X | X | X |
| L43 | T57 | New | X | X | X | X |
| R45 | V55 | New | X | X | X | X |
| Q46 | H19 | | | | New | X |
| E53 | R51 | | | New | X | X |
| Y54 | L69 | | New | X | X | X |
| V55 | R45 | New | X | X | X | X |
| L56 | L67 | New | X | X | X | X |
| T57 | L43 | | New | X | X | X |
| V58 | K65 | New | X | X | X | X |
| N59 | V41 | | | New | X | X |
| F60 | K63 | New | X | X | X | X |
| K63 | F60 | | New | X | X | |
| K65 | F58 | New | X | X | X | X |
| L67 | L56 | New | X | X | X | X |
| L69 | Y54 | New | X | X | X | X |
| S70 | R78 | | | New | X | X |
| N72 | Q76 | | | New | X | X |
| G75 | N72 | | | New | X | X |
| C77 | F84 | | | | New | X |
| R78 | S70 | | | New | X | |
| V79 | L82 | | New | X | X | X |
| L82 | V79 | | New | X | X | X |
| F84 | C77 | | | | New | X |
| D89 | S86 | | | | New | |
| M90 | S86 | | New | X | X | X |
| L91 | I87 | New | X | X | X | X |
| E92 | F88 | New | X | X | X | X |
| H93 | D89 | New | X | X | X | X |
| F94 | M90 | New | X | X | | X |
| R95 | E92 | New | X | X | X | X |
| V96 | E92 | New | X | X | X | X |
| H97 | H93 | New | X | X | X | X |
| I99 | V109 | New | X | X | X | X |
| G104 | L101 | | | | New | X |
| V109 | S107 | | | | New | X |
| V112 | G40 | New | x | X | X | X |
| V115 | F42 | | | | New | X |

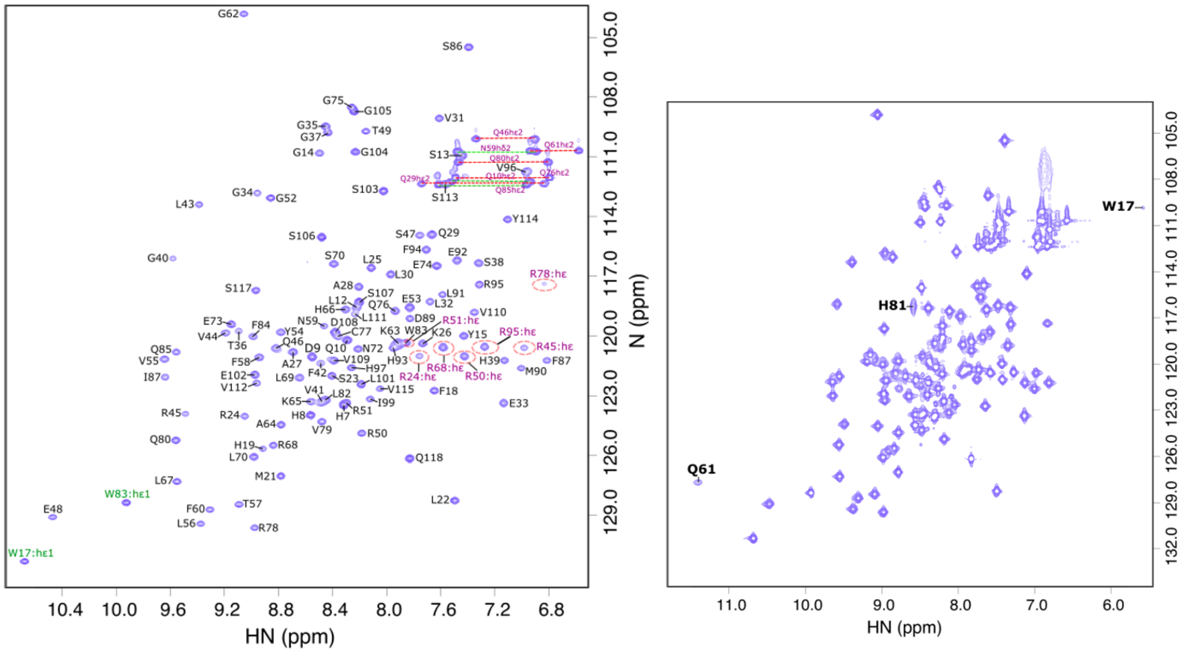


**Figure S1.** $^{1}H$,$^{15}N$ HSQC spectrum of SH2 recorded at pH 6 and 298 K. Left, $^{1}H$ $^{15}N$ HSQC shows the backbone resonances of SH2 indicated by sequence number and residue name. Red circled peaks and green labelled peaks illustrate the sidechain NHε of Arg and NHε of Trp, respectively, which were assigned by NOESY spectra. The top right corner contains the sidechain amide (Nε-Hε2 of Gln, Nδ-Hδ2 of Asn) peaks, which are connected by horizontal lines and assigned based on NOESY assignment. Right, $^{1}H$ $^{15}N$ HSQC showing the low intensity peaks (W17, Q61, H81).

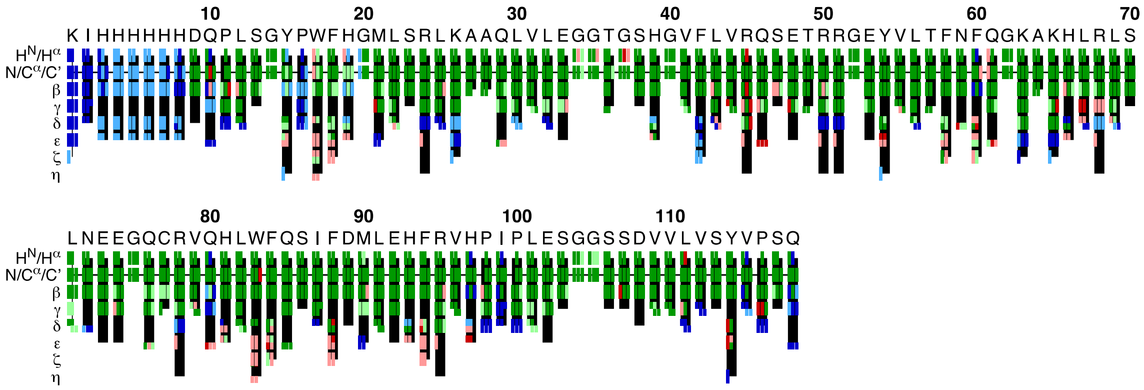


**Figure S2**. An overview graphic of CYANA assignment for SH2 residues 1-118, using manual assignment as a reference. On the top are the residue and residue name. In general, light colours indicate less confident assignments and dark colours illustrate strong assignments. Blue color indicates no reference assignment available; green color shows that the assignment by CYANA agreed with the reference assignment within defined tolerance value; red color shows the assignments that do not agree with the reference assignment; black color represents the assignments that were found in reference assignment but not assigned by CYANA. For side-chain assignments, in the center is the heavy atom assignment with hydrogen atoms to the left and right. In branched side-chains, the row is divided into upper and lower parts.

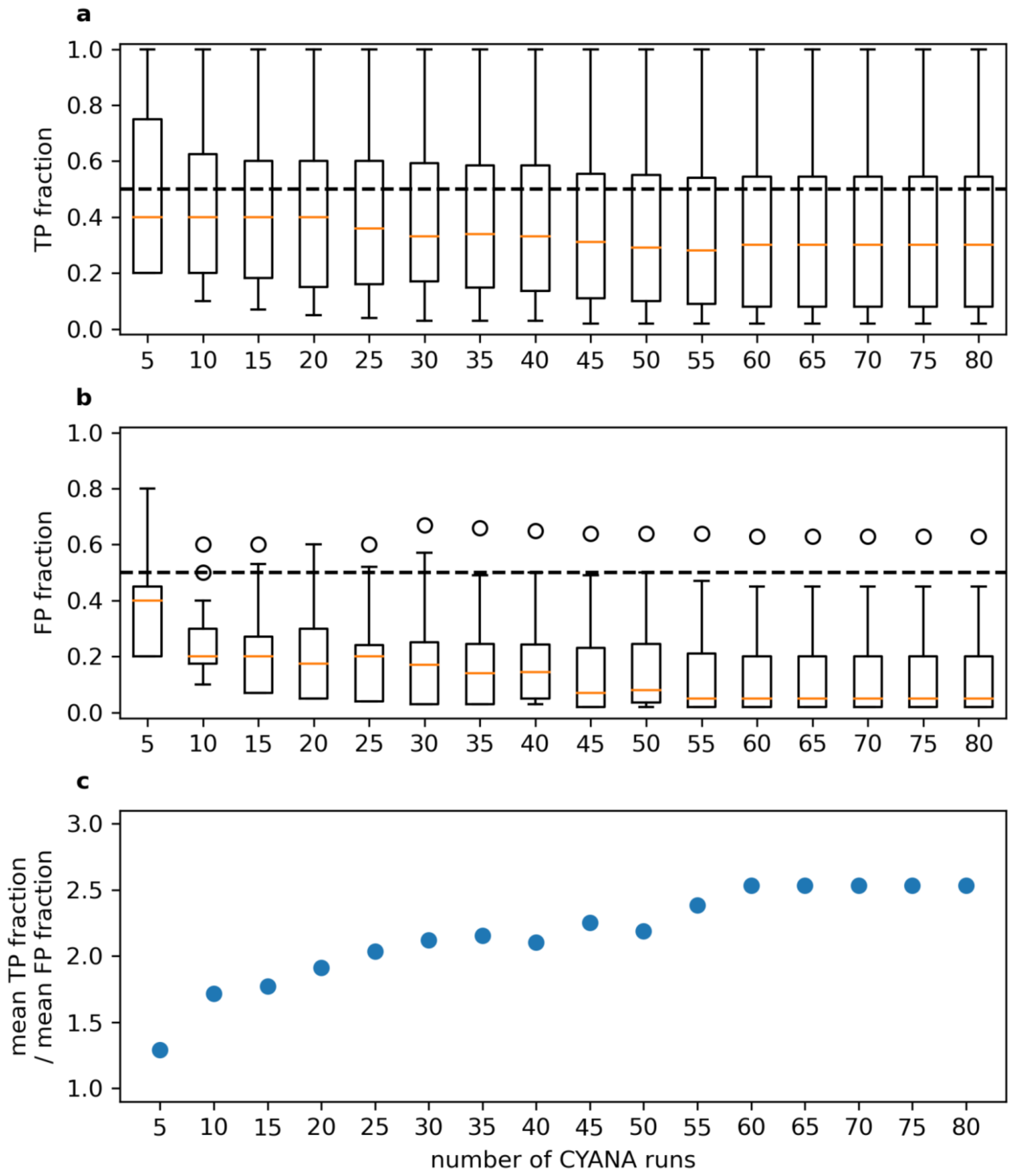


**Figure S3**. **Determination of the optimum number of repeated calculations**. a) Fraction of hydrogen bonds that appear in NMR ensembles that are also present in the X-ray structure PDB 5w3r (true positives [TP]) plotted against the number of CYANA runs performed (each run generated an ensemble of 20 models). b) Fraction of hydrogen bonds that appear in NMR models that are not present in the X-ray structure (false positives [FP]) plotted against the number of CYANA runs. c) The ratio of the mean fraction of true positives to the mean fraction of false negatives plotted against the number of CYANA runs. Both the fraction of true positives and true negatives decrease as more CYANA runs are performed. False positives decay at a faster rate so that increasing the number of runs increases the chance that a true positive is identified. After 60 runs the ratio of true positives to false negatives stabilises.

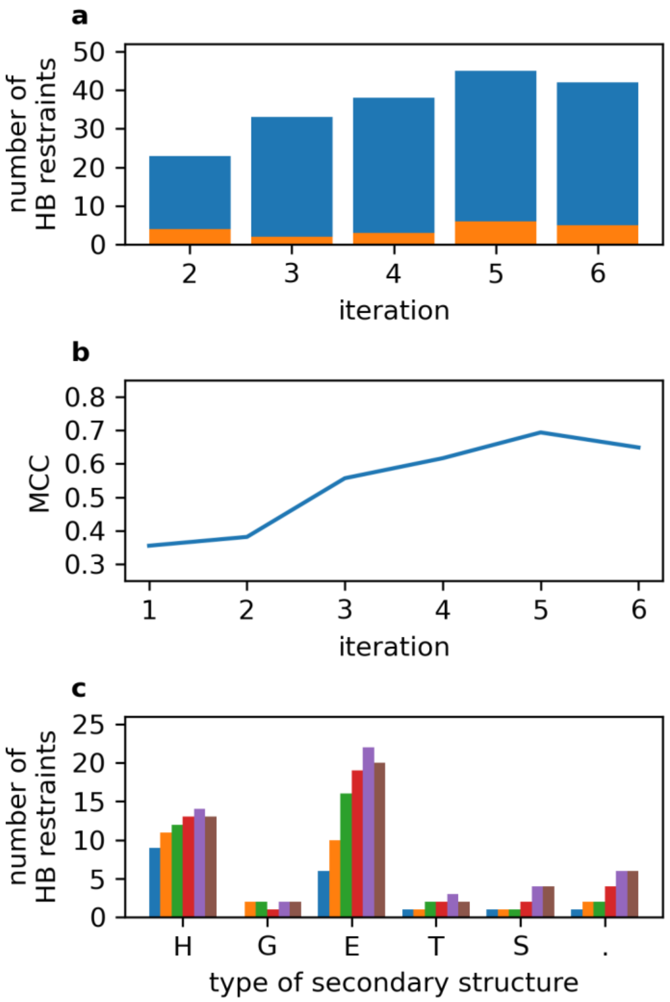


**Figure S4.** A comparison of hydrogen bond restraints (HBRs) throughout the 6 iterations to those seen in the crystal structure (PDB ID 5w3r). (a) Total number of HBRs at each iteration, colored blue if seen in the crystal structure or orange if not. (b) Matthew's correlation coefficient for HBR correctness, defined as

$$MCC = \frac{(TP \times TN) - (FP \times FN)}{\sqrt{(TP + FP)(TP + FN)(TN + FP)(TN + FN)}}$$

A true positive (TP) is an amide with a hydrogen bond restraint that is also hydrogen bonded in the X-ray structure; a false positive (FP) is an amide with a hydrogen bond restraint but is not hydrogen bonded in the X-ray structure; a true negative (TN) is an amide which does not have a hydrogen bond restraint and is not hydrogen bonded in the X-ray structure; and a false negative (FN) is an amide which does not have a hydrogen bond restraint but is hydrogen bonded in the X-ray structure. Categorising the amides this way allows us to compute the Matthew's correlation coefficient (MCC) which returns a value between −1 and +1. A coefficient of +1 represents a perfect agreement, 0 no better than random and −1 total disagreement. (c) Total number of HBRs for different categories of structure, for each of the 6 iterations. (H) Helix, (G) $3_{10}$ helix (E) β-sheet regions, (T) turns, (S) bends and (.) regions with no regular secondary structure.

```
                        10        20        30        40        50        60
SH2B1|mouse|    KIHHHHHHDQPLSGYPWFHGMLSRLKAAQLVLEGGTGSHGVFLVRQSETRRGEYVLTFNF
SH2B1|homo |    ----GSHMDQPLSGYPWFHGMLSRLKAAQLVLTGGTGSHGVFLVRQSETRRGEYVLTFNF
                       520       530       540       550       560       570

                        70        80        90       100       110
SH2B1|mouse|    QGKAKHLRLSLNEEGQCRVQHLWFQSIFDMLEHFRVHPIPLESGGSSDVVLVSYVPSQ  →(118)
SH2B1|homo |    QGKAKHLRLSLNAAGQCRVQHLHFQSIFDMLEHFRVHPIPLESGGSSDVVLVSYVPSS  →(628)
                       580       590       600       610       620
```

**Figure S5.** Sequences of the SH2 domains of mouse (this work) and human (crystal structure) SH2B1, including the N-terminal tags used for the two constructs. Changes in amino acid sequence are highlighted; the mutations E583A, E584A and W591H were introduced to improve crystallization and are not present in the native human sequence (McKercher et al., 2018). The mutations had little effect on peptide affinity and so are presumed to have little effect on the structure.

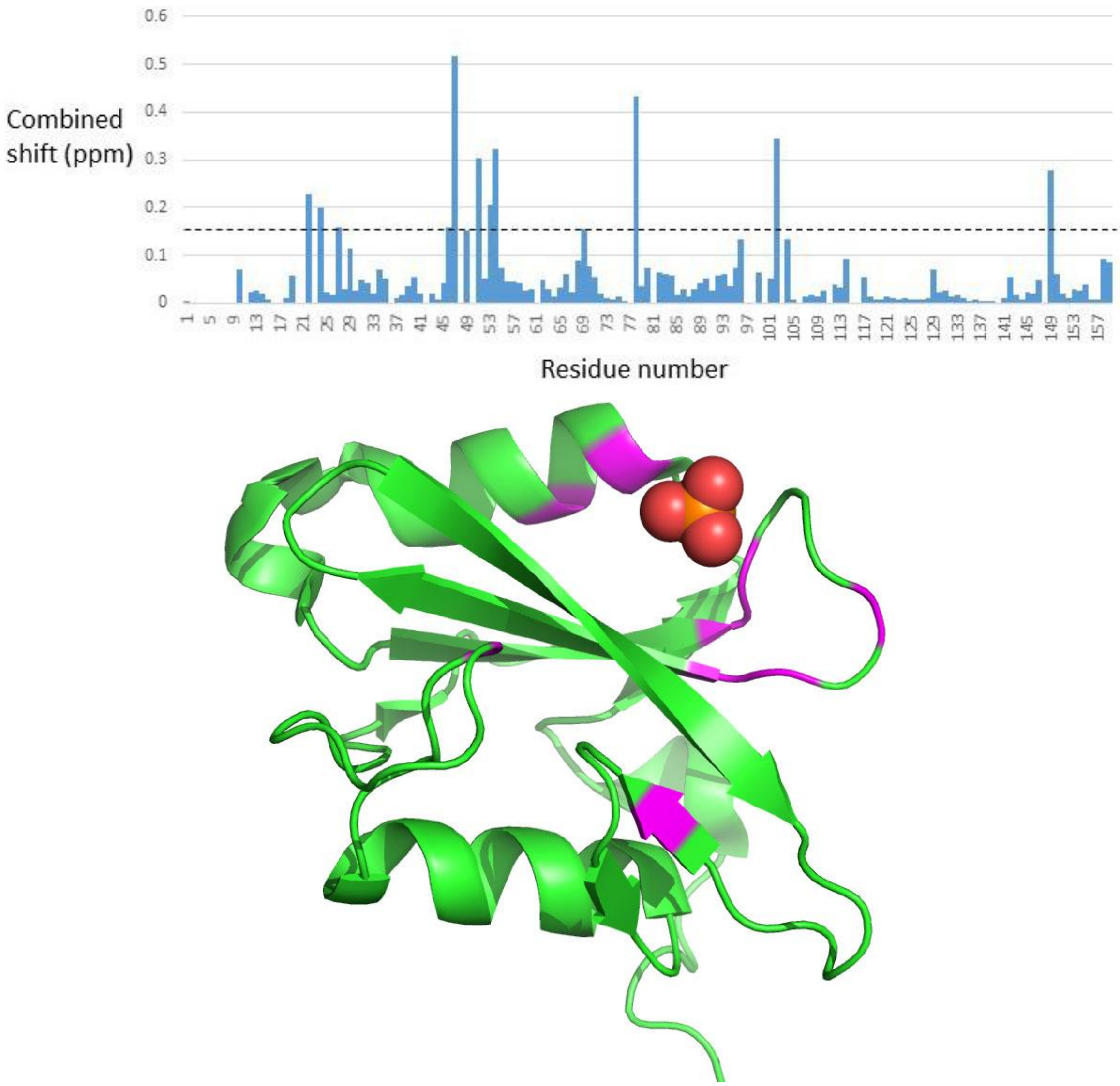


**Figure S6.** (top) Chemical shift changes on changing the buffer from 50 mM Tris to 50 mM phosphate, pH 6. The changes are the weighted change in $H_N$ and N shifts, calculated as $\sqrt{(\delta H^2 + (0.14\delta N)^2)/2}$ (Williamson, 2013). The dashed line is at 2 x (standard deviation). (bottom) Crystal structure of SH2 from SH21B (PDB ID 5w3r), showing the bound phosphate ion. Residues with chemical shift changes > 2xSD on addition of phosphate are coloured in magenta. The 46-51 loop is just below and to the right of the phosphate ion.

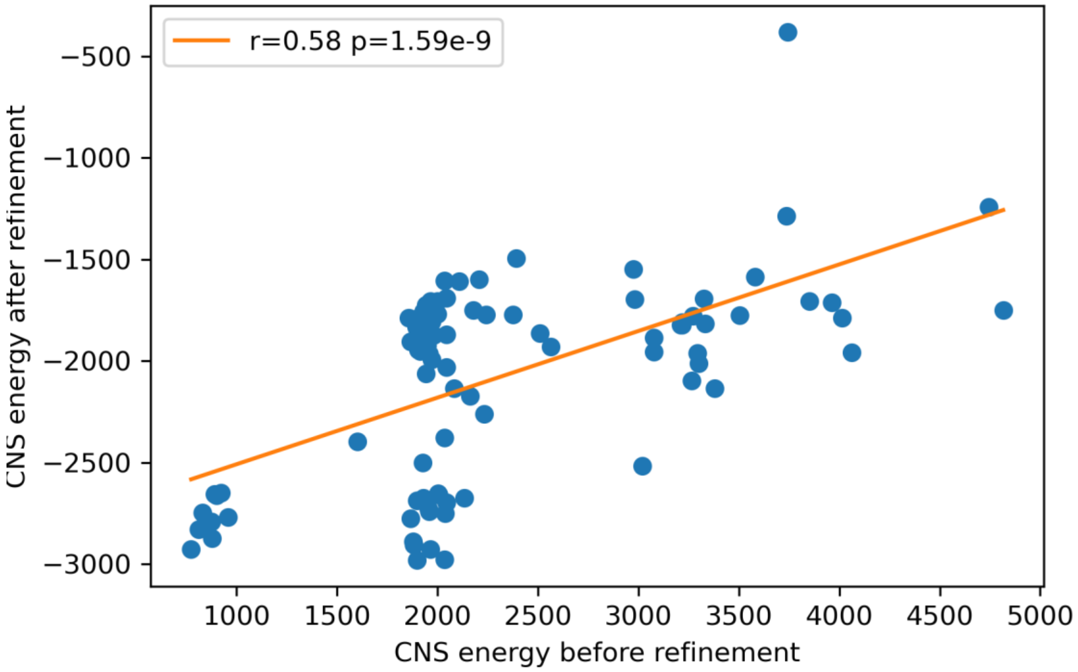


**Figure S7.** Relationship between ensemble energies before and after refinement in water. 100 structures were calculated using the simplified initial forcefield. They were then refined in explicit solvent. Of the two sets of 20 lowest energy structures before and after refinement, there are 12 structures found in both sets. Two of the 100 structures are not shown here, which have much higher energies in both sets.

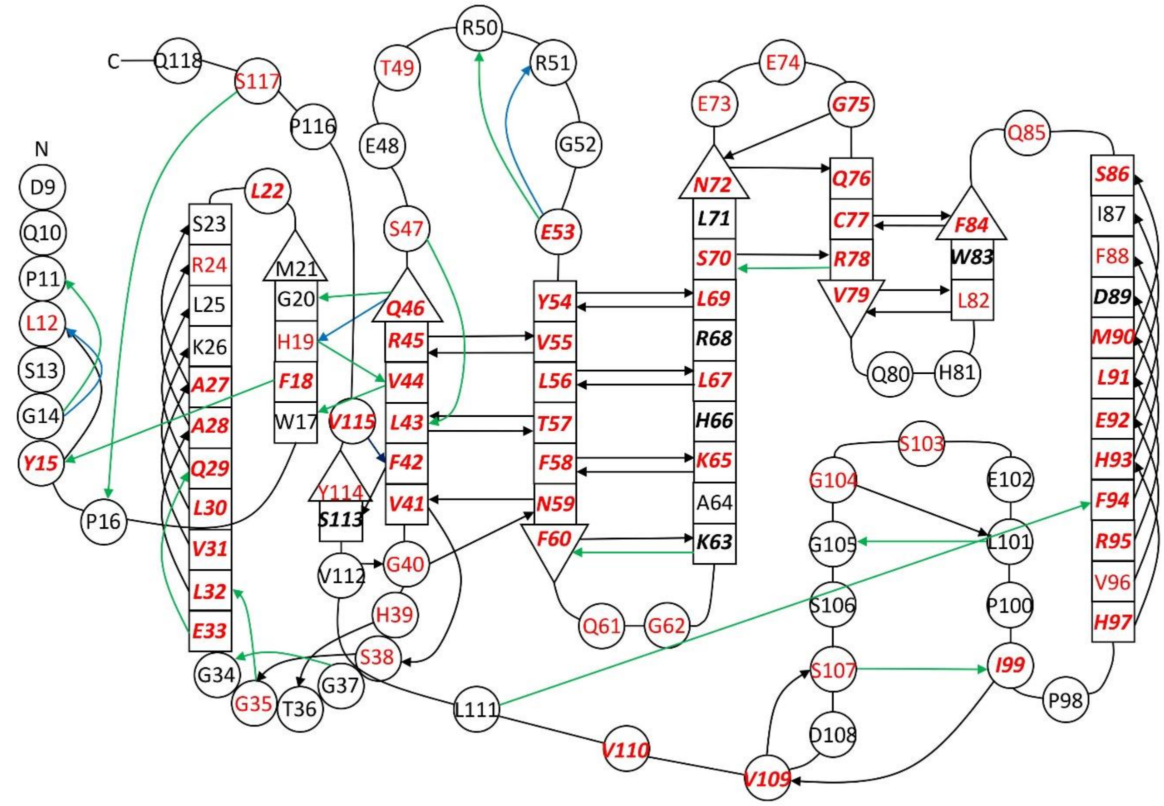


**Figure S8.** A comparison of hydrogen bonds identified in this study with those in the crystal structure 5w3r. Residue colouring is the same as Figure 3. Hydrogen bonds are indicated by arrows going from the NH to CO. Bonds in black are in both structures; bonds in green are in the crystal structure only, and bonds in blue are in the NMR structure only.
